# Bi-allelic variants in *WDR47* lead to neuronal loss causing a rare neurodevelopmental syndrome with corpus callosum dysgenesis in humans

**DOI:** 10.1101/2023.12.22.572779

**Authors:** Efil Bayam, Peggy Tilly, Stephan C. Collins, José Rivera Alvarez, Meghna Kannan, Lucile Tonneau, Bruno Rinaldi, Romain Lecat, Noémie Schwaller, Sateesh Maddirevula, Fabiola Monteiro, João Paulo Kitajima, Fernando Kok, Mitsuhiro Kato, Ahlam A. A. Hamed, Mustafa A. Salih, Saeed Al Tala, Mais Hashem, Hiroko Tada, Hirotomo Saitsu, Sylvie Friant, Zafer Yüksel, Mitsuko Nakashima, Fowzan S. Alkuraya, Binnaz Yalcin, Juliette D. Godin

## Abstract

The corpus callosum (CC) is the largest interhemispheric connection that is largely formed by the axons of layer 2/3 callosal projection neurons (CPNs) through a series of tightly regulated cellular events, including neuronal specification, migration, axon extension and branching. Defects in any of those steps may prevent the proper development of the corpus callosum resulting in a spectrum of disorders collectively referred to as corpus callosum dysgenesis (CCD). Here, we report four unrelated families carrying bi-allelic variants in *WDR47* presenting with CCD together with other neuroanatomical phenotypes such as microcephaly, cerebellar abnormalities and hydrocephalus. Using a combination of *in vitro* and *in vivo* mouse models and complementation assays, we show that independently from its previously identified functions in neuronal migration and axonal extension, WDR47 is required for survival of callosal neurons by contributing to the maintenance of mitochondrial and microtubule homeostasis. We further provide evidence that severity of the CCD phenotype is determined by the degree of the loss of function caused by the human variants. Taken together, we identify *WDR47* as a causative gene of a new neurodevelopmental syndrome characterized by corpus callosum abnormalities and other neuroanatomical malformations.

## INTRODUCTION

The corpus callosum consists of 190 million axons in an adult human brain making it the largest commissure in mammals. This interhemispheric connection is largely formed by the axons of layer 2/3 callosal projection neurons (CPNs) through a series of tightly regulated cellular events involving commitment of newborn neurons to CPN identity, migration to the proper cortical layer, axonal extension and guidance towards the midline with the aid of secreted and contact-mediated axon-guidance cues ^1, 2,3^. Defects in any of those steps may prevent the proper development of the corpus callosum resulting in a spectrum of disorders collectively referred to as corpus callosum dysgenesis (CCD). The phenotypes associated with CCD are heterogeneous and can range from agenesis of the corpus callosum (AgCC) with complete or partial loss of the CC to an unusually thin (hypoplasia) and thick (hyperplasia) CC. CCD can be found either isolated which has generally a more favorable outcome or together with other neuroanatomical phenotypes (NAPs ^4^) such as microcephaly presenting a higher risk of cognitive and neuropsychiatric disorders ^2, 5^. To date, there are more than 300 known genetic syndromes associated with CCD, however the majority remain undiagnosed at the molecular level and their underlying pathophysiological mechanisms under characterized ^5^.

WDR47 is a poorly-studied microtubule associated protein particularly enriched in neurons in humans ^6^ and mice ^7, 8^ that we previously identified as a novel regulator of mouse CC development using a large-scale knock-out (KO) screen ^4^. *Wdr47* is essential for mouse survival and brain morphogenesis in a dose dependent manner ^7, 9^. Total loss of *Wdr47* leads to lethality immediately after birth (*Wdr47*^tm1b/tm1b^). Yet, a fraction (nearly 6%) of mice expressing 30% of *Wdr47* (*Wdr47*^tm1a/tm1a^) survive and display severe neuroanatomical abnormalities including primary microcephaly and severe loss of commissural fibers, including CC, that manifest as hyperactivity and sensory motor gating abnormalities ^7^. At the cellular level, WDR47 has been shown to regulate the proliferation of late progenitors and neuronal migration *in vivo* in mouse developing brains as well as axonal and dendritic development *in vitro* in primary culture of neurons ^7, 9, 10^. While WDR47 is known to regulate cellular processes that are critical for proper brain development such as microtubule stability ^9, 10^, intracellular transport ^11^ and autophagy ^7^, it is still unclear whether these regulatory functions are required for proper development of callosal projection neurons.

Despite the clear role of Wdr47 in mouse brain development and function, there is yet no clear association of *WDR47* variants with neurological disorders in humans. The only exception is a genome-wide association study (GWAS) that identified *WDR47* as a hub gene in Alzheimer’s disease ^12^. Here, we provide the first evidence of a causal relationship between variants in *WDR47* and human neuroanatomical phenotypes. We report four bi-allelic loss-of-function variants in patients presenting with severe brain malformations including CCD, microcephaly, thin cortex, hydrocephalus and cerebellar atrophy. Using a combination of *in vitro* and *in vivo* mouse models and complementation assays, we show that Wdr47 is required for survival of callosal neurons. Although independent from its previously identified roles in neuronal migration and axonal extension, WDR47’s role in neuronal survival partly relies on its cell-autonomous function on microtubule cytoskeleton. We further demonstrate that the degree of the loss of function imposed by the different human *WDR47* bi-allelic variants varies and dictates the severity of the CC phenotype. Taken together, our data imply *WDR47* as an important gene for neuroanatomical disorders in humans whose screening should be considered in unexplained syndromic cases involving corpus callosum dysgenesis, microcephaly and other neuroanatomical phenotypes.

## RESULTS

### Human *WDR47* bi-allelic missense variants delineate severe neurodevelopmental phenotypes with corpus callosum dysgenesis (CCD), microcephaly, intellectual disability and epilepsy

We identified, through the GeneMatcher platform^13^ and data sharing, a series of five cases (M01 to M05; five males; M01 and M02 are siblings) from four unrelated families worldwide (Families 1 to 4 living in Sudan, Japan, Saudi Arabia and Brazil, respectively) with bi-allelic missense variants (**Figures 1A-D**).

**Figure 1:**
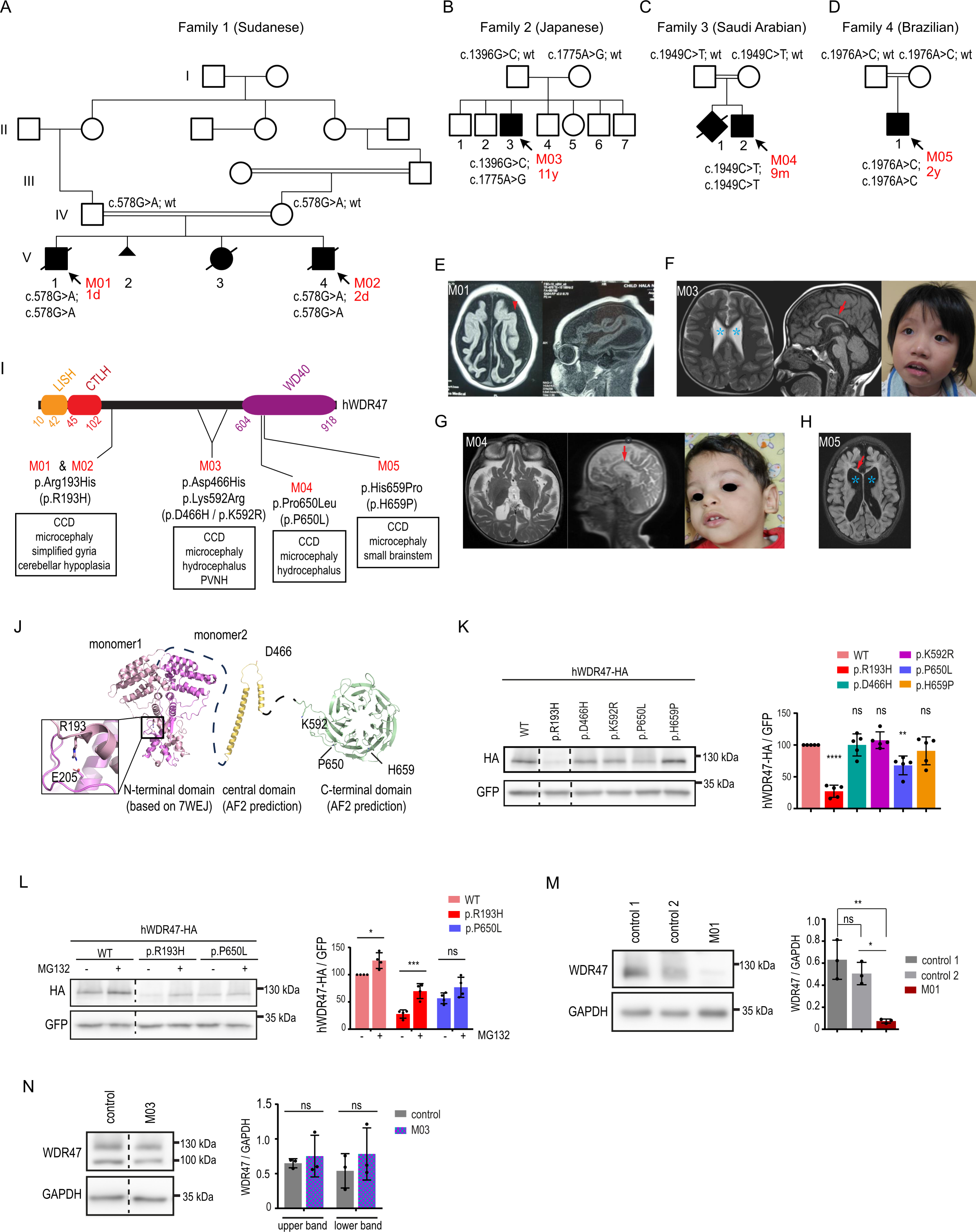
Clinical and brain imaging data from five male patients with *WDR47* recessive loss-of- function variants. (A-D) Pedigrees showing the segregation of the identified *WDR47* variants with the syndrome in consanguineous families (exception of Family 2). **(E-H)** Axial and sagittal T1 and T2- weighted brain MRI and facial features of patients with *WDR47* variants. The red arrowhead indicates the simplified gyral pattern of the cortex, the blue stars show ventricular dilatations, and the red arrows point to thin corpus callosum. **(I)** Schematic representation of the human WDR47 (hWDR47) protein depicting the LISH, CTLH and WD40 domains and the position of the mutated amino acids for M01&M02 (p.(Arg193His) (R193H)), M03 (p.(Asp466His)/p.(Lys592Arg) (D466H/L592R)), M04 (p.(Pro650Leu) (P650L)) and M05 (p.(His659Pro) (H659P)). **(J)** Structural modeling of the human WDR47 (hWDR47) protein with an Alphafold2-derived atomic model. Homodimerization of the N-terminal featuring the LISH and CTLH domains is shown based on the crystal structure of mouse WDR47-NTD intertwined dimer. The position of the mutated amino acids for M01 & M02 (p.R193), M03 (p.D466) and (p.K592), M04 (p.P650) and M05 (p.H659) and E205 residue that makes a salt bridge with the p.R193 are indicated. **(K)** Western blot analysis of extracts from N2A cells transfected with the indicated HA tagged *h*WDR47 constructs showed variable effect of variants on WDR47 protein levels. GFP was used as a transfection control. Dotted lines indicate where the membrane was cut. Data (means ± sd) from 5 independent cultures was analyzed by one-way ANOVA, with Bonferroni’s multiple comparison test. **(L)** MG132 treatment rescued the decreased levels of p.R193H variant suggesting proteasomal degradation of the mutant protein. Data (means ± sd) from 4 independent cultures was analyzed by two-way ANOVA, with Bonferroni’s multiple comparison test. **(M-N)** Western blot analysis showing that endogenous levels of WDR47 is **(M)** decreased in fibroblast extracts from patient M01 and **(N)** unchanged in lymphoblastoid cell extracts from patient M03. **(M)** Fibroblast and **(N)** lymphoblastoid cell lines from two and one healthy subjects were used respectively as controls. Data (means ± sd) from 3 cultures was analyzed by **(M)** one- way ANOVA, with Bonferroni’s multiple comparison test and **(N)** unpaired t-test. ns, non-significant; *P < 0.05; **P < 0.005; ***P < 0.0005; ****P < 0.0001. Dotted lines indicate the position where the membrane was cut.

Variants were inherited from healthy consanguineous parents in an autosomal recessive manner at the exception of the Japanese Family 2 that showed a compound heterozygous mode of inheritance where the parents are healthy non-consanguineous. The median age at the time of the last evaluation was 2,75 years (ranging from 1 day old to 11 years old).

The clinical features are summarized in **Table 1**. All cases had neuroanatomical phenotypes (NAPs) with the most frequent phenotypes being corpus callosum dysgenesis (5/5) and microcephaly (5/5), followed by a reduced size of the hindbrain (3/5), hydrocephalus (2/5), simplified gyral pattern (2/5) and periventricular nodular heterotopia (1/5) (**Figures 1E-H**). Other features included mild to severe intellectual disability (5/5), epilepsy (5/5), developmental delays (2/3), muscle tone defects (2/3), and craniofacial features (2/2). Craniofacial characteristics in case M03 included widely spaced eyes, epicanthus and a short philtrum. In case M04, the features were hypertelorism, medial flaring of eyebrows, thick upper and lower lips, elevated ear lobules and a low auricle. Both cases, M03 and M04, showed a tented upper lip (**Figures 1F-G**).

**Table 1:**
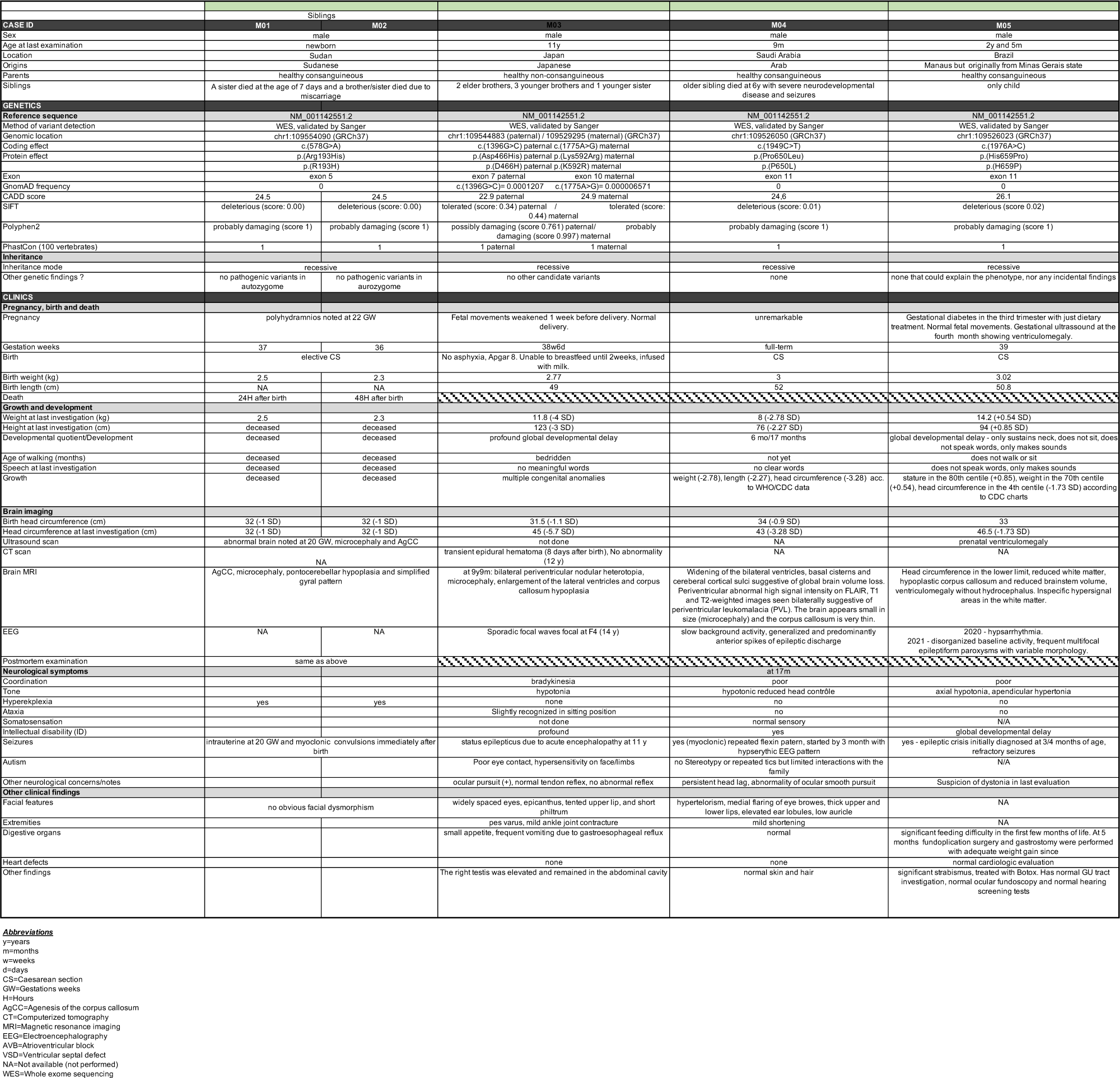

We identified four unique bi-allelic missense *WDR47* variants that occur within conserved residues located in the N-terminal domain (NTD ^14^) (NM_001142551.2, c.(578G>A), p.(Arg193His), M01 & M02), the central region (NM_001142551.2, c.(1396G>C), p.(Asp466His) paternal, c.(1775A>G), p.(Lys592Arg) maternal, M03), and the C-terminal WD40 domain (NM_001142551.2, c.(1949C>T), p.(Pro650Leu), M04 and c.(1976A>C), p.(His659Pro), M05) (**Figure 1I and Supplementary** Figures 1A-D). Various predictions scores, such as CADD, SIFT, Polyphen2 and PhastCon, indicated deleteriousness at conserved residues (**Table 1**). None of the five variants are reported in public databases, including dbSNP, 1000 genomes and gnomAD at the homozygous state. To evaluate the impact of identified variants in the full-length WDR47, we employed structural modeling with an Alphafold2-derived atomic model. The variants within the central domain (p.(Asp466His)) or close or within the C-terminal WD40 (p.(Lys592Arg), p.(Pro650Leu), p.(His659Pro)), localize to solvent-exposed regions and, although their specific effects remain uncertain, are unlikely to disrupt the correct fold of the protein (**Figure 1J**). In contrast, the p.(Arg193His) substitution, that maps to WDR47-NTD involved in homodimerization ^14^, likely interferes with the formation of the intertwined WDR47-NTD dimer. This disruption is likely due to the alteration of the salt bridge that Arginine 193 forms with Glutamate 205, a corresponding residue in the other monomer (**Figure 1J**). This weakened WDR47 homodimerization interface was also confirmed with the PISA server (http://www.ebi.ac.uk/pdbe/prot_int/pistart.html) ^15^.

To interrogate the molecular effect of the *WDR47* variants, we first assessed WDR47 protein expression levels after overexpression of HA-tagged WT and mutant constructs in N2A neuroblastoma cell line. The variants have various effect on protein expression level ranging from no effect for p.(Asp466His), p.(Lys592Arg) and p.(His659Pro) variants, to moderate (-32%, p.(Pro650Leu)) and severe (-72%, p.(Arg193His)) decrease compared to the wild-type (WT) human protein (**Figure 1K**). Decreased expression level of Arg193His mutant protein was rescued after treatment with the proteasome inhibitor MG132, suggesting instability of the WDR47 mutant Arg193His protein (**Figure 1L**). Such rescue was not observed with the Pro650Leu, indicating that the Pro650Leu WDR47 protein is not targeted for proteasomal degradation (**Figure 1L**). Next, we examined *WDR47* protein levels in patient’s samples when available (M01 and M03). Though the levels of *WDR47* transcripts remained stable (**Supplementary** Figure 1E), immunoblotting revealed a complete loss of p.(Arg193His) WDR47 protein levels in fibroblasts derived from M01 (**Figure 1M**), further indicating that the mutant Arg193His WDR47 protein is unstable. Western-Blot analysis in lymphoblastoid cell line obtained from the patient M03 carrying the compound heterozygous variant (p.(Asp466His), p.(Lys592Arg)) confirmed the absence of phenotype at the protein expression level (**Figure 1N**). Overall, we identified five missense substitutions in the *WDR47* gene, that differently impact on WDR47 protein abundance, in five patients presenting with severe neurodevelopmental delay associated with corpus callosum dysgenesis (CCD), microcephaly and other NAPs.

### Neuron-specific *Wdr47* knock-out in mice recapitulates NAPs observed in human patients

Interestingly, the clinical manifestations of the patients are recapitulated in previously studied constitutive *Wdr47* KO mouse models ^7, 9, 16^ that display severe neuroanatomical phenotypes including CCD and microcephaly among other NAPs (**Tables 1-2, Supplementary** Figure 2A). This indicates that *Wdr47*- deficient mice are valuable models to understand the pathogenicity of *WDR47* loss-of-function variants in humans. As NAPs could arise from defects in progenitors and neurons or both, we investigated the extent to which NAPs induced by *WDR47* deficiency are intrinsic to neurons. We first conditionally deleted *Wdr47* specifically in early glutamatergic projection neurons in the dorsal telencephalon using the neuronal basic helix-loop-helix Nex^Cre^ mouse line ^17^ **(Supplementary** Figure 2A). Loss of *Wdr47* in E18.5 Nex^Cre^;*Wdr47*^fl/fl^ embryos was validated by immunoblotting **(Supplementary** Figure 2B ). To ensure high reproducibility, we acquired neuroanatomical data in both Nex^Cre^;*Wdr47*^fl/fl^ and *Wdr47*^tm1a/tm1a^ simultaneously using our recently established neuroanatomical quantification approach for embryonic brains ^18^. Our histology pipeline was designed based on two coronal sections at Bregma +2.19mm and Bregma +3.51mm, spanning 17 developmentally distinct brain regions for 32 quantifiable brain parameters of area, height and width measurements (**Figures 2A-D and Supplementary Table 1**). Each brain image was carefully checked for quality control before analysis taking into consideration the correct stereotaxic position and the symmetry both along the dorsoventral and rostrocaudal orientations. In line with our previous report ^7^, we identified NAPs in *Wdr47*^tm1a/tm1a^ constitutive E18.5 KO embryonic brains (**Figure 2A, Supplementary** Figure 2A) including CCD (-24%, *p*=0.04), hypoplasia of the hippocampus, thinning of the motor cortex, and reduced size of the internal capsule (ic) and anterior part of the anterior commissure (aca) (**Figure 2B and Table 2**). However, unlike in *Wdr47*^tm1a/tm1a^ constitutive E18.5 KO, Nex^Cre^;*Wdr47*^fl/fl^ embryonic brains did not reveal any change in the CC area, width or height despite the strong reduction in the size of other commissures including the internal capsule (-27%, *p*=0.0019) and the anterior commissure (-80%, *p*=4.56E-07) (**Figures 2C-D**, **Table 2**). This suggests that the 24% decrease in the size of the CC observed in the constitutive KO mice stem from defects likely involving non-neuronal cells (such as glial cells or radial glia progenitors) at early stages of CC development. Similar to constitutive KO mice ^7^ and a previous report ^9^, Nex^Cre^;*Wdr47*^fl/fl^ pups died soon after birth confirming that the neonatal death of the *Wdr47*-deficient mice is attributable to brain defects.

**Table 2:**
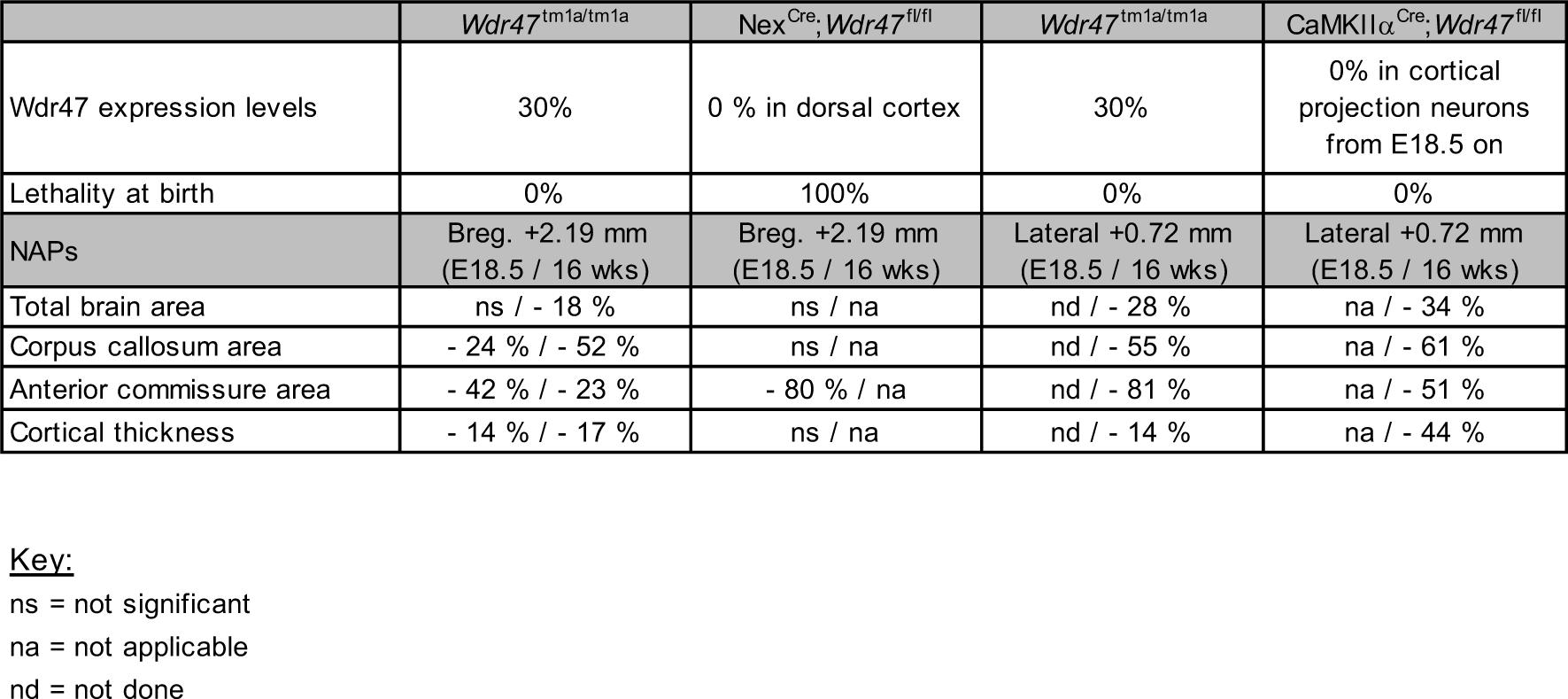

**Figure 2:**
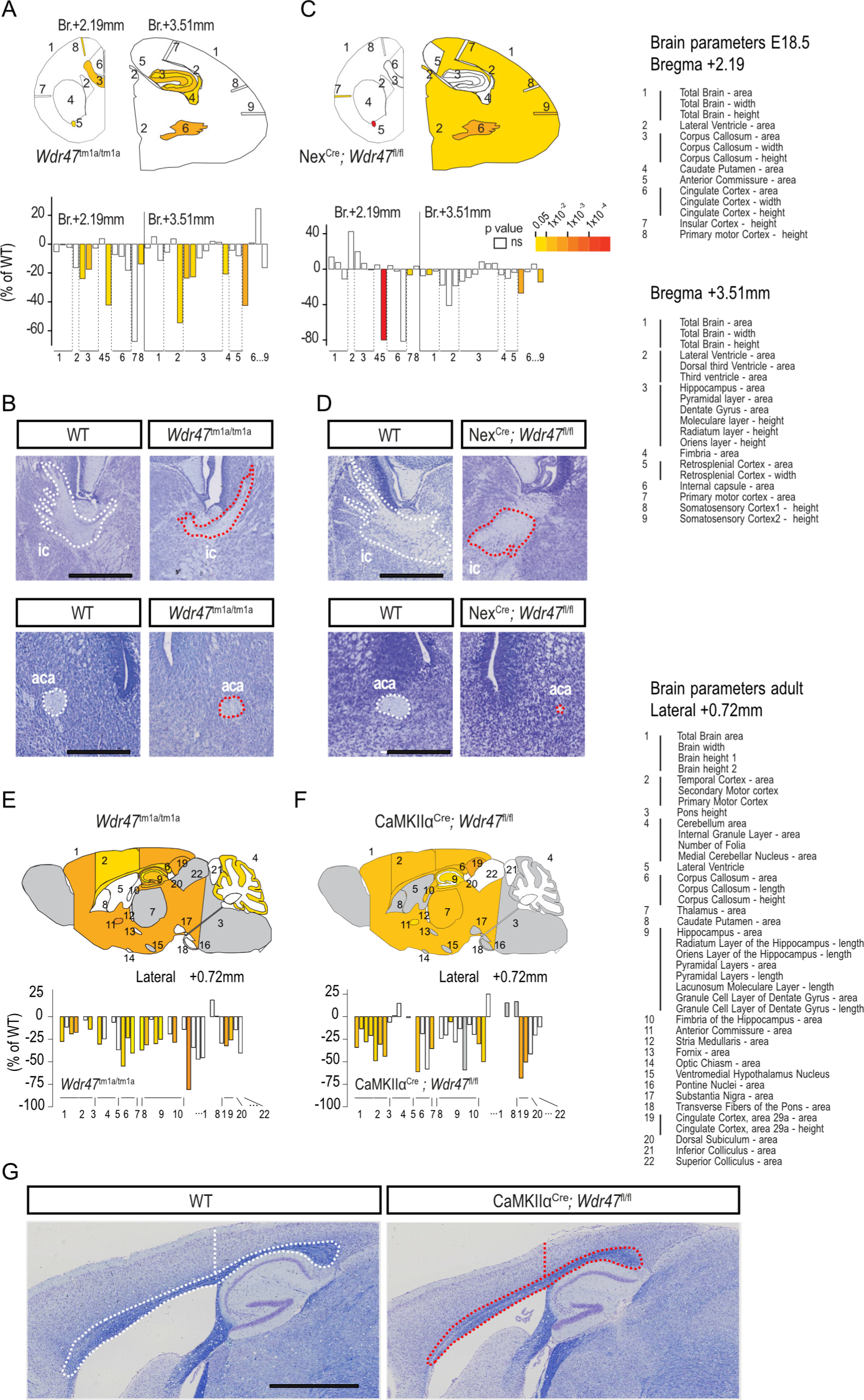
Neuroanatomical studies conducted in various mouse models reveal the neuronal origin of the observed defects. (**A**) *Top*. Schematic representation of the 17 brain regions assessed at Bregma +2.19 mm and +3.51 mm in E18.5 *Wdr47^tm1a/tm1a^*mice (n=5 WT *vs* n=4 *Wdr47*^tm1a/tm1a^). Colored regions indicate the presence of at least one significant parameter within the brain region at the 0.05 level. White indicates a p-value > 0.05, grey shows not enough data to calculate a p-value. *Bottom.* Histograms of percentage change relative to matched WT animals (set as 0) for each of the measured parameters (listed in **Supplementary Table 1** and on the right-hand side of the Figure). (**B**) Representative brain images stained with Nissl of embryonic WT and *Wdr47*^tm1a/tm1a^ mice showing the internal capsule (ic) at Bregma +3.51 mm and the anterior part of the anterior commissure (aca) at Bregma +2.19 mm. (**C**) *Top*. Schematic representation of the 17 brain regions assessed at Bregma +2.19 mm and +3.51 mm in E18.5 Nex^Cre^;*Wdr47*^fl/fl^ mice (n=6 WT *vs* n=6 cKO). Colored regions indicate the presence of at least one significant parameter within the brain region at the 0.05 level. White indicates a p-value > 0.05, grey shows not enough data to calculate a p-value. *Bottom.* Barplots of percentage change relative to matched WT animals (set as 0) for each of the measured parameters. (**D**) Representative brain images stained with Nissl of embryonic WT and Nex^Cre^;*Wdr47*^fl/fl^ embryonic mice showing the internal capsule (ic) at Bregma +3.51 mm and the anterior part of the anterior commissure (aca) at Bregma +2.19 mm. (**E**) *Top.* Schematic representation of the 22 brain regions quantified at Lateral +0.72 mm on parasagittal section from adult *Wdr47*^tm1a/tm1a^ mice, aged 16 weeks mice (n=3 WT *vs* n=3 *Wdr47*^tm1a/tm1a^). Colored regions indicate the presence of at least one significant parameter within the brain region at the 0.05 level. White indicates a p-value > 0.05, grey shows not enough data to calculate a p-value. *Bottom.* Histograms of percentage change relative to WT mice (set as 0) for each of the measured parameters (listed in **Supplementary Table 1** and on the right-hand side of the Figure). (**F**) *Top.* Schematic representation of the 22 brain regions quantified at Lateral +0.72 mm on parasagittal section from adult CaMKIIα^Cre^;*Wdr47*^fl/fl^ at 16 weeks of age (n=3 WT *vs* n=2 cKO). Colored regions indicate the presence of at least one significant parameter within the brain region at the 0.05 level. White indicates a p-value > 0.05, grey shows not enough data to calculate a p-value. *Bottom.* Histograms of percentage change relative to WT mice (set as 0) for each of the measured parameters. (**G**) Representative brain images stained with Nissl-luxol of adult WT (left) and CaMKIIα^Cre^;*Wdr47*^fl/fl^ (right) mice showing the area of the corpus callosum and the height of the cortex at Lateral +0.72 mm.

The neonatal death of the Nex^Cre^;*Wdr47*^fl/fl^ pups did not allow to study the neuron-specific involvement of *Wdr47* at later stages of CC development. To circumvent this problem, we used the calcium/calmodulin-dependent protein kinase II alpha subunit CaMKIIα^Cre^ line ^19^ to delete *Wdr47* specifically in post-mitotic neurons in the forebrain starting from E18.5 ^20^, when the peak of *Wdr47* expression is occurring (https://apps.kaessmannlab.org/evodevoapp/) ^6^ (**Supplementary** Figure 2A). With respect to the 3Rs (replacement, reduction and refinement) principles to limit animal use, we studied brain anatomy using one time point in adult CaMKIIα^Cre^; *Wdr47*^fl/fl^ (16 weeks of age) as the mice were viable. RT-qPCR analysis confirmed the decreased expression of *Wdr47* transcripts in adult cortices (**Supplementary** Figure 2C). Using our established pipeline for the neuroanatomical quantification of 22 brain structures in the adult mouse totaling 40 measurements at Lateral +0.72mm ^21^, we identified a similar profile of NAPs in CaMKIIα^Cre^; *Wdr47*^fl/fl^ and *Wdr47*^tm1a/tm1a^ adult mice with the exception of the cerebellar anomalies which were absent in the CaMKIIα^Cre^; *Wdr47*^fl/fl^ mice (**Figures 2E-F**). The corpus callosum was smaller in size by 61% (*p*=0.008) and 55% (*p*=0.027) and the total brain area by 34% (*p*=0.0095) and 28% (*p*=0.014) in CaMKIIα^Cre^; *Wdr47*^fl/fl^ and *Wdr47*^tm1a/tm1a^ adult mice, respectively **(Figure 2G and Table 2)**. These combined findings in conditional and constitutive mouse models showed that, although non-neuronal cells likely contribute to prenatal CC abnormalities, there is a major involvement of post-mitotic neurons to neuroanatomical phenotypes in young adult (**Table 2**).

### *Wdr47* deficiency leads to perinatal loss of callosal interhemispheric connections by impairing neuronal survival

Considering the presence of CCD in all the patients (**Table 1**) and severe CC anomalies in *Wdr47*- deficient mice (**Table 2**), we next focused on the development of callosal projection neurons (CPNs) that extend axons toward the midline that *in fine* innervate the contralateral cortex through extensive branching. We performed acute depletion of *Wdr47* specifically in callosal neurons using *in utero* electroporation (IUE) of vectors allowing the expression of Cre-GFP or GFP under the control of the NeuroD promoter together with the pCAG2-mScarlet reporter plasmid in *Wdr47*^fl/fl^ or *Wdr47*^fl/WT^ embryonic cortices at E15.5, when the CPNs are born. At postnatal day 2 (P2), soon after the axons cross the midline, the proportion of neurons able to project their axon was decreased upon *Wdr47* depletion as indicated by the reduced density (-55%) of scarlet-positive axons in the white matter (WM) of *Wdr47*^fl/fl^ (IUE NeuroD-Cre-GFP) cortices compared to controls (NeuroD-GFP in *Wdr4*7^fl/fl^ or *Wdr47*^fl/WT^) (**Figures 3A-C**). In addition, almost one third of the *Wdr47*-deficient axons failed to reach the midline (31% of axons arriving to the midline in controls compared to 11% in *Wdr47*^fl/fl^ pups after IUE with Cre) (**Figures 3A-B and 3D**). However, once they reached the midline, axons depleted for *Wdr47* showed similar ability to cross the midline than control axons (**Figures 3A-B and 3E**). Of note, no aberrant axonal projection or midline crossing were observed in *Wdr47*^fl/WT^ pups electroporated with NeuroD-Cre-GFP (**Figures 3A-E**) suggesting that *Wdr47* is haplosufficient for axon extension. Strikingly, at P4, while axons extended further into the contralateral hemisphere in control and haploinsufficient (NeuroD-Cre-GFP in *Wdr47*^fl/WT^) condition, the proportion of scarlet-positive axons at the midline was severely decreased upon *Wdr47* deletion (IUE Neuro-Cre-GFP in *Wdr47*^fl/fl^) (**Figure 3F**). A closer examination of *Wdr47*-deficient neurons in the upper cortical plate (uCP) revealed aberrant cellular appearance with loss of bipolar organization and fragmented neurites (**Supplementary** Figure 3A) as well as presence of several pyknotic nuclei and apoptotic cells (positive for activated Caspase 3) at P3 and P4 but not at P2 (**Supplementary** Figures 3A and 3B). As a consequence, at P8, most of the *Wdr47*-depeleted neurons have disappeared and the remaining neurons failed to maintain their bipolar morphology and axonal projections as shown by the absence of scarlet-positive axons at the midline (**Figure 3F**, insets). Altogether, these results suggested that WDR47 has a pro-survival function in callosal neurons at perinatal stages.

**Figure 3:**
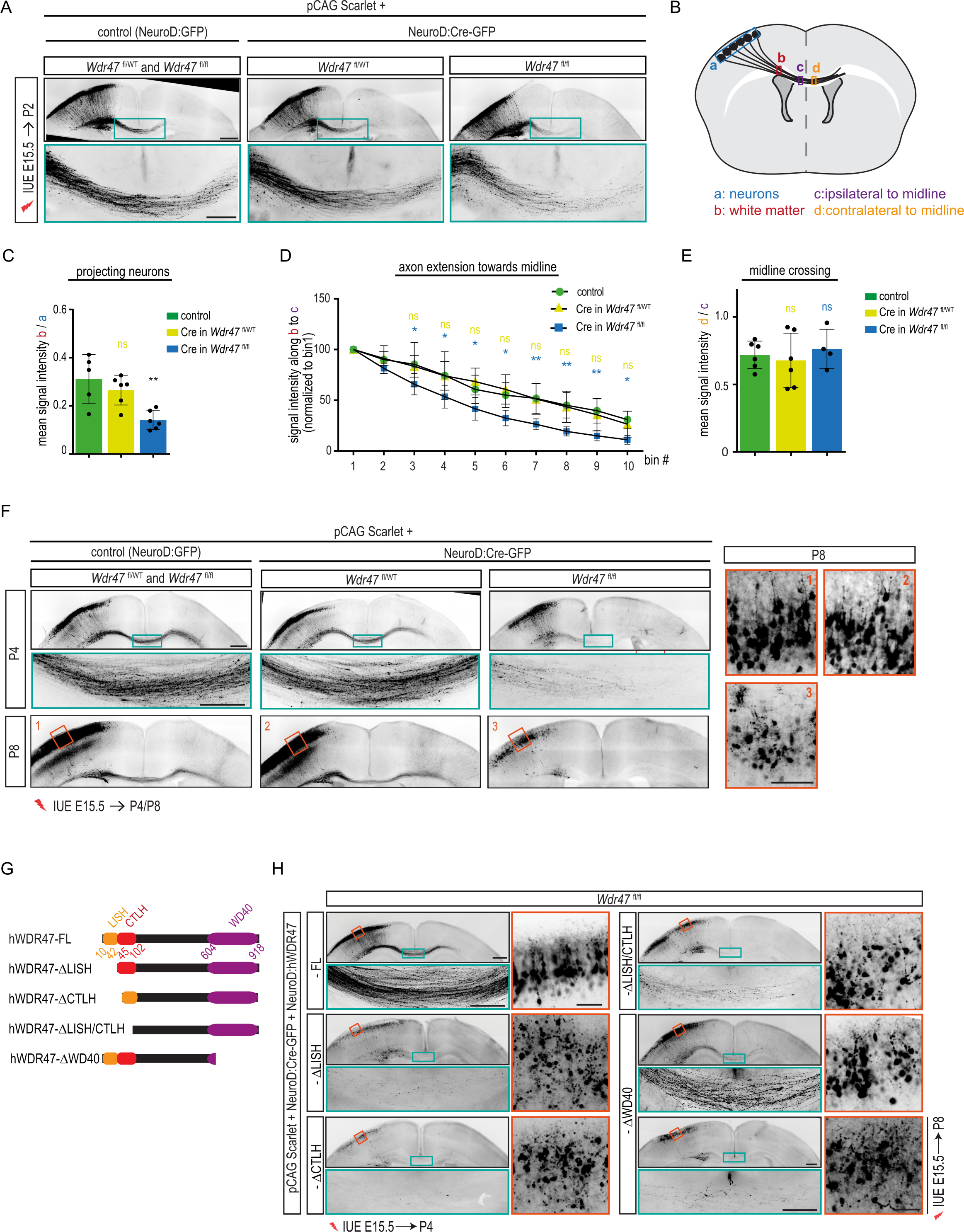
Deletion of *Wdr47* impedes interhemispheric connectivity through impaired neuronal survival. (A) Coronal sections of P2 mouse brains electroporated at E15.5 with pCAG:Scarlet plasmid together with either a control (NeuroD:Ires:GFP) or a NeuroD:Cre-GFP vector. Scarlet positive electroporated neurons are depicted in black. Close-up views of the green boxed area show axon extension defects in *Wdr47*^fl/fl^ pups but not in *Wdr47*^fl/WT^ pups or control conditions. Scale bars: 500µm and 200µm (green boxed inset). **(B)** Schematic describing the methods used to quantify **(C)** the percentage of projecting neurons (mean intensity of the scarlet signal in the white matter (red box) normalized on the mean intensity in the cortical plate (blue box)) **(D)** axon extension towards midline (intensity was plotted along a line that was divided into 10 equal bins from the immediate start of CC (red box) till the midline (purple box); intensity in bin1 was considered 100% and intensity in each bin was normalized to bin1) and **(E)** midline crossing (mean intensity of the scarlet signal just after the midline (yellow box) is normalized on the mean intensity just before midline crossing (purple box)). **(C-E)** Quantification of **(C)** projecting neurons **(D)** axon extension towards midline **(E)** midline crossing. Data (means ± sd) from at least 4 pups from 2-3 different litters per condition were analyzed by **(C and E)** one- way ANOVA, with Bonferroni’s multiple comparison test, **(D)** two-way ANOVA, with Bonferroni’s multiple comparison test. **(F)** Coronal sections of P4 and P8 mouse brains electroporated at E15.5 with pCAG:Scarlet plasmid together with either a control (NeuroD:Ires:GFP) or a NeuroD:Cre-GFP vector. Scarlet positive electroporated neurons are depicted in black. Close-up views of the green and orange boxed area show loss of CC at P4 and loss of neuronal morphology at P8 respectively in *Wdr47*^fl/fl^ pups but not in *Wdr47*^fl/WT^ pups or control conditions. At least 4 pups from 2-3 different litters per condition was analyzed. Scale bars: 500µm, 200µm (blue boxed inset) and 100µm (red boxed inset). **(G)** Schematic representation of the WDR47 protein depicting different domains and the truncated Wdr47 constructs used in **(H)** to rescue the phenotype induced by the loss of *Wdr47.* **(H)** Coronal sections of P4 and P8 mouse brains electroporated at E15.5 with pCAG:Scarlet plasmid and NeuroD:Cre-GFP vector together with a truncated hWDR47 construct. Scarlet positive electroporated neurons are depicted in black. Close-up views of the green and orange boxed area show that at P4 ΔLISH, ΔCTLH and ΔLISH/CTLH fail to rescue the loss of CC and neuronal morphology while ΔWD40 construct could partially restore the survival defects. Scale bars: 500µm, 200µm (blue boxed inset) and 50µm (red boxed inset). ns, non- significant, *P < 0.05, **P < 0.005.

WDR47 is a multidomain protein with N-terminally located LISH and CTLH domains and seven C-terminally located WD40 domains (**Figure 3G**). Therefore, to study which of those domains could mediate the pro-survival function of WDR47, we tested the ability of truncated WDR47 constructs to rescue the phenotype induced by the loss of *Wdr47.* We performed IUE of NeuroD-Cre-GFP and pCAG2-mScarlet plasmids together with either a NeuroD-driven full length (hWDR47-FL) or truncated human WDR47 construct lacking different domains in E15.5 *Wdr47*^fl/fl^ or *Wdr47*^fl/WT^ embryos and analyzed the scarlet-positive neurons and their axons at P4. In accordance with a cell autonomous function of Wdr47, introduction of WDR47-FL construct rescued the survival defects (**Figure 3H**). Expression of all N-terminal deletion proteins (hWDR47-τιLISH, -τιCTLH and -τιLISH/CTLH) failed to rescue CC phenotype, as assessed by the absence of Scarlet positive axons in the midline (**Figure 3H**). Although *Wdr47*-deficient neurons expressing truncated WDR47 proteins that lack the WD40 domain (WDR47-ΔWD40) were able to project their axons toward the contralateral cortex as the WDR47-FL expressing neurons, they started displaying signs of degeneration at P4 (**Figure 3H**, insets). At P8, most of the axons at the midline disappeared indicating that expression of WDR47-ΔWD40 likely delays the onset of the CC phenotype (**Figure 3H**). Of note, 10 times less of WD40 plasmid was used to reach similar expression level between all the constructs (**Supplementary** Figure 3C). Given the absence of phenotypes after IUE of Cre in *Wdr47*^fl/WT^ pups at P4 (**Figure 3F and Supplementary** Figures 3A and 3B), we used them as controls and showed that none of those constructs impair neuronal survival in control condition (**Supplementary** Figure 3D). Altogether these results indicate that WDR47 is required cell autonomously to protect CPNs from sudden neuronal death *in vivo* and that both N- and C-terminal domains of Wdr47 are indispensable for its pro-survival function.

### Loss of *Wdr47* causes neuronal death independently of its role in migration and axonal extension

As Wdr47 was previously shown to regulate radial migration ^7, 9^, we further investigated whether faulty migration at early stages could drive cell death of callosal neurons at later stages. First, we investigated whether the roles of Wdr47 in cell survival and neuronal migration involve the same domains. We tested for the restoration of migration phenotype induced by the loss of *Wdr47* ^7, 9^ by introducing FL or truncated hWDR47 constructs together with NeuroD-Cre-GFP using IUE in E14.5 *Wdr47*^fl/fl^ embryos and quantified the distribution of GFP-positive neurons in the cortical plate 4 days later. As we previously did not observe any difference in migration between control and *Wdr47* haploinsufficient neurons ^7^, we used IUE of NeuroD-Cre-GFP in *Wdr47^f^*^l/WT^ embryos as control. Consistent with previous reports ^7, 9^, acute depletion of *Wdr47* resulted in severe migration defects with 54% of cell reaching the upper cortical plate (uCP) compared to 77% in control condition (**Figures 4A and 4B**). While introduction of FL-hWDR47 construct fully restored the migration (73% GFP+ cells in uCP), none of the N-terminal truncated constructs could rescue the phenotype (53%, 48%, 51%, GFP+ cells in the uCP for the WDR47-ΔLISH, -ΔCTLH and -ΔLISH/CTLH constructs respectively) (**Figures 4A and 4B**). At the opposite, Wdr47 protein lacking the C-terminal WD40 domain partially rescued the migration phenotype (70% GFP+ cells in uCP) (**Figures 4A and 4B**). Of note, expression of none of those constructs affected migration in *Wdr47*^fl/WT^ embryos (**Supplementary** Figures 4A and 4B). These results suggested that N-terminally located LISH and CTLH domains likely mediate functions of WDR47 in both neuronal migration and survival whereas WD40 repeats are at least partially dispensable for migration.

**Figure 4:**
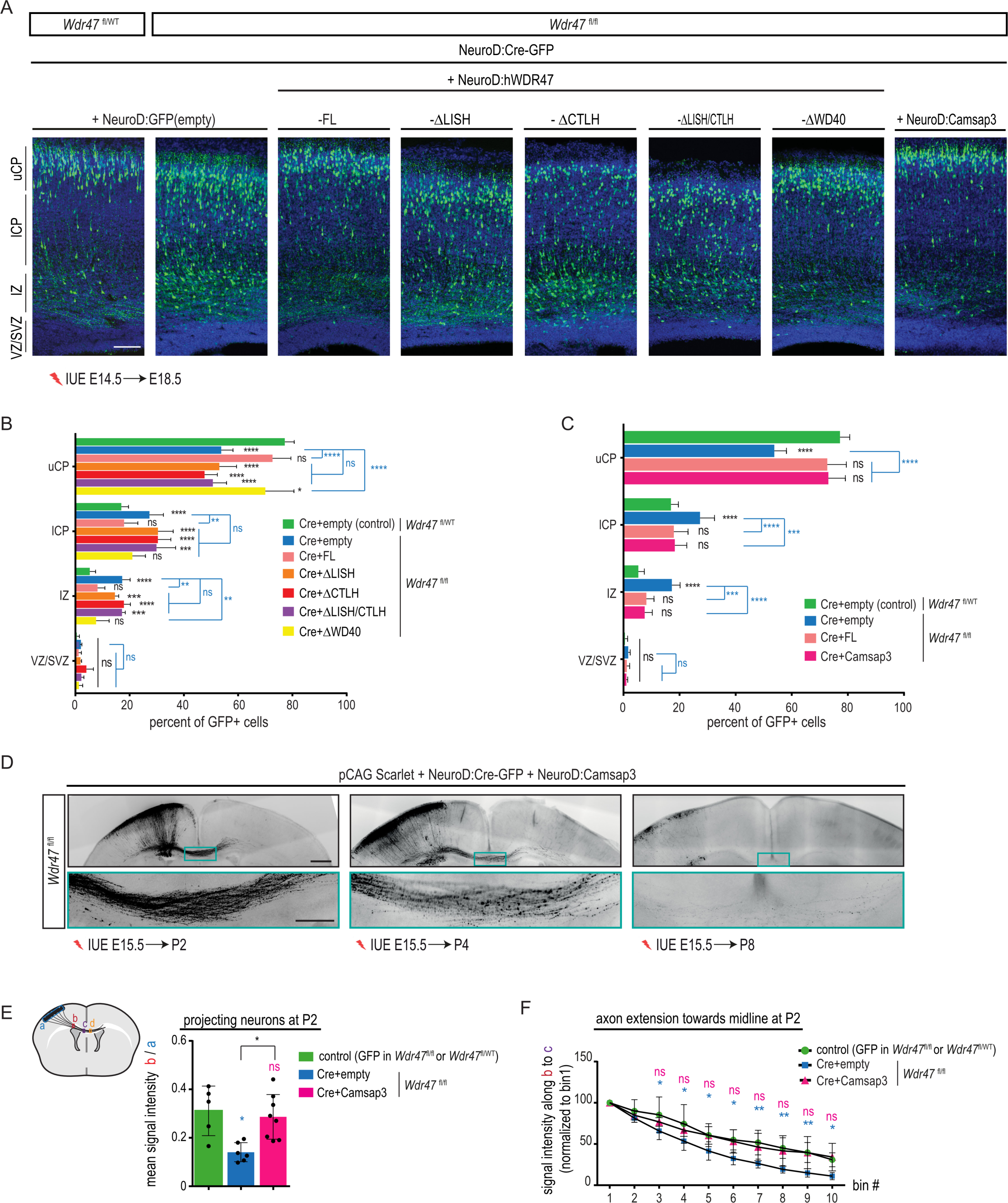
Function of WDR47 in neuronal survival is independent from its function in neuronal migration and axon extension. (A) Coronal sections of E18.5 mouse cortices, four days after *in utero* electroporation with NeuroD-Cre-GFP together with either NeuroD:empty vector or a different rescue vector (either NeuroD:hWDR47-FL or a truncated NeuroD:hWDR47 construct or NeuroD:Camsap3). GFP positive electroporated cells are depicted in green. Nuclei are stained with DAPI. Scale bar: 100µm. **(B-C)** Analysis of the percentage of electroporated GFP-cells in different regions (uCP, lCP, IZ and VZ /SVZ) showing effect of introducing **(B)** different truncated Wdr47 constructs and **(C)** Camsap3 on neuronal migration. Data (means ± s.d) from at least three embryos from 2-3 different litters per condition was analyzed by two-way ANOVA, with Bonferroni’s multiple comparisons test. uCP, Upper cortical plate; lCP, Lower cortical plate; IZ, intermediate zone; VZ, ventricular zone; SVZ, subventricular zone. **(D)** Coronal sections of P2, P4 and P8 mouse brains electroporated at E15.5 with pCAG:Scarlet, NeuroD:Cre-GFP and NeuroD:Camsap3 plasmids. Scarlet positive electroporated neurons are depicted in black. Close-up views of the green boxed area show that axon extension defects are rescued upon introduction of Camsap3 at P2 however CC is still lost at P8. Scale bars: 500µm and 200µm (green boxed inset) **(E-F)** Quantification of **(E)** projecting neurons **(F)** axon extension towards midline. Data (means ± sd) from at least 5 pups per condition were analyzed by **(E)** one-way ANOVA, with Bonferroni’s multiple comparison test, **(F)** two-way ANOVA, with Bonferroni’s multiple comparison test. ns, non-significant; *P < 0.05; **P < 0.005; ***P < 0.001; ****P < 0.0001.

Next, as Camsap3, a microtubule (MT) minus end binding protein that recruits WDR47 to MT minus ends, had been shown to rescue the migration and neuronal polarization defects of *Wdr47*-deficient neurons ^9^, we tested whether rescuing the migration and axonal extension defects by introducing Camsap3 in *Wdr47*-deficient neurons would also rescue neuronal survival defects. We expressed mouse Camsap3 together with NeuroD-Cre-GFP in *Wdr47*^fl/fl^ embryos by IUE at E14.5 and analyzed the distribution of GFP+ cells at E18.5. In parallel, we performed IUE at E15.5 and assessed axonal projection at the midline at P2 and neuronal survival at P4 and P8. Interestingly, although overexpression of Camsap3 fully restored the migration defects at E18.5 (**Figures 4A and 4C**) ^9^ as well as axon extension defects at P2 and P4 (**Figures 4D-4F**), it only slightly delayed neuronal death as shown by the absence of CC and severe degeneration of neurons at P8 (**Figure 4D**). Moreover, the analysis of CC of Camsap3 adult KO mice revealed a thinner CC, likely due to the specific loss of medial fibers as no notable differences were observed in the lateral fibers connecting the somatosensory cortex (**Supplementary** Figure 4C **and Supplementary Table 1**). This is in contrast with the agenesis of the corpus callosum observed in *Wdr47*^tm1a/tm1a^ KO mice ^7^, supporting that only some of the WDR47-related functions during brain development are mediated by Camsap3.

To sum up, our results showed that although they rely on the same domains, pro-survival function of WDR47 is independent of its roles in migration and axonal growth and involves downstream effectors other than Camsap3.

### Wdr47 protects callosal neurons from apoptosis

We next sought to understand the mode of cell death induced upon loss of *Wdr47*. As we previously reported a negative correlation between WDR47 expression and activation of autophagy ^7^, we tested whether autophagic cell death could cause the survival defects induced by loss of *Wdr47*. We turned to the yeast model and addressed first the requirement of LISH/CTLH domains for yeast growth as a proxy of survival. While expression of human WDR47-FL allowed normal growth ^7^, expression of hWDR47-ΔLISH or -ΔCTLH constructs strongly impaired growth in WT yeast (**Supplementary** Figures 5A-C), reminiscent of their requirement for the survival of cortical neurons. Then, we investigated the involvement of those domains in autophagy inducing conditions upon nitrogen starvation, by following the yeast Atg8 (homologue to mammalian LC3) reporter. In line with a role of WDR47 in inhibiting autophagy ^7^, mCherry-Atg8 did not accumulate in the lumen of the vacuole and instead remained cytoplasmic upon expression of full length WDR47 (**Supplementary** Figure 5D). Surprisingly, expression of hWDR47-ΔLISH and -ΔCTLH inhibited accumulation of mCherry-Atg8 in the vacuolar lumen similarly to the FL construct suggesting that those 2 domains, that are indispensable for the function of WDR47 in survival of cortical neurons (**Figure 3H**) and yeast (**Supplementary** Figures 5B-C), are not required for inhibiting autophagy (**Supplementary** Figure 5D). According to an autophagy independent mechanism, expression of Wdr47-ΔLISH and -ΔCTLH constructs in the Δoxa1 yeast strain that is deficient in respiration and that lacks mitochondrial DNA ^22^ did not lead to any growth defects (**Supplementary** Figures 5B-C) suggesting that growth defects induced by those constructs likely involve mitochondrial- rather than autophagy-dependent mechanisms.

To further identify which mode of cell death is triggered upon *Wdr47*-deficiency, we used a simplified *in vitro* system that allows to study dynamics of neuronal death at the single cell resolution. We transfected primary neuronal cultures established from cortices of E15.5 *Wdr47*^WT/WT^ and constitutive *Wdr47*^tm1b/tm1b^ knock-out embryos (hereafter named WT and HOM, respectively) with a pCAG2-mScarlet plasmid.

Monitoring of individual scarlet-positive neuron from day *in vitro* (DIV) 3 to DIV10 by time lapse imaging (**Figure 5A**), revealed a burst of neuronal death starting at DIV8 in HOM cultures leading to a decrease by half of the number of surviving neurons in HOM cultures compared to WT at DIV10 (**Figures 5A- C**).Treatment of HOM primary neurons with Qvd-Oph, a pan-Caspase inhibitor, at DIV2, fully rescued the death of *Wdr47* knock-out neurons at DIV10 (62% survival in Qvd-Oph-treated HOM cultures compared to 69% survival in WT cultures) (**Figures 5B-C**), whereas treatment with Necrostatin and Ferrostatin, necroptosis and ferroptosis inhibitors respectively, did not have any effect on survival (34%, 36% and 49% survival in untreated, necrostatin- and ferrostatin-treated HOM cultures respectively) (**Figures 5B- C**). Of note, none of the drugs affected survival of WT neurons (**Supplementary** Figures 5E-F). Taken together, these results demonstrated that loss of *Wdr47* leads to a severe, Caspase dependent, early-onset burst of neuronal death that manifests as CC dysgenesis *in vivo*.

**Figure 5:**
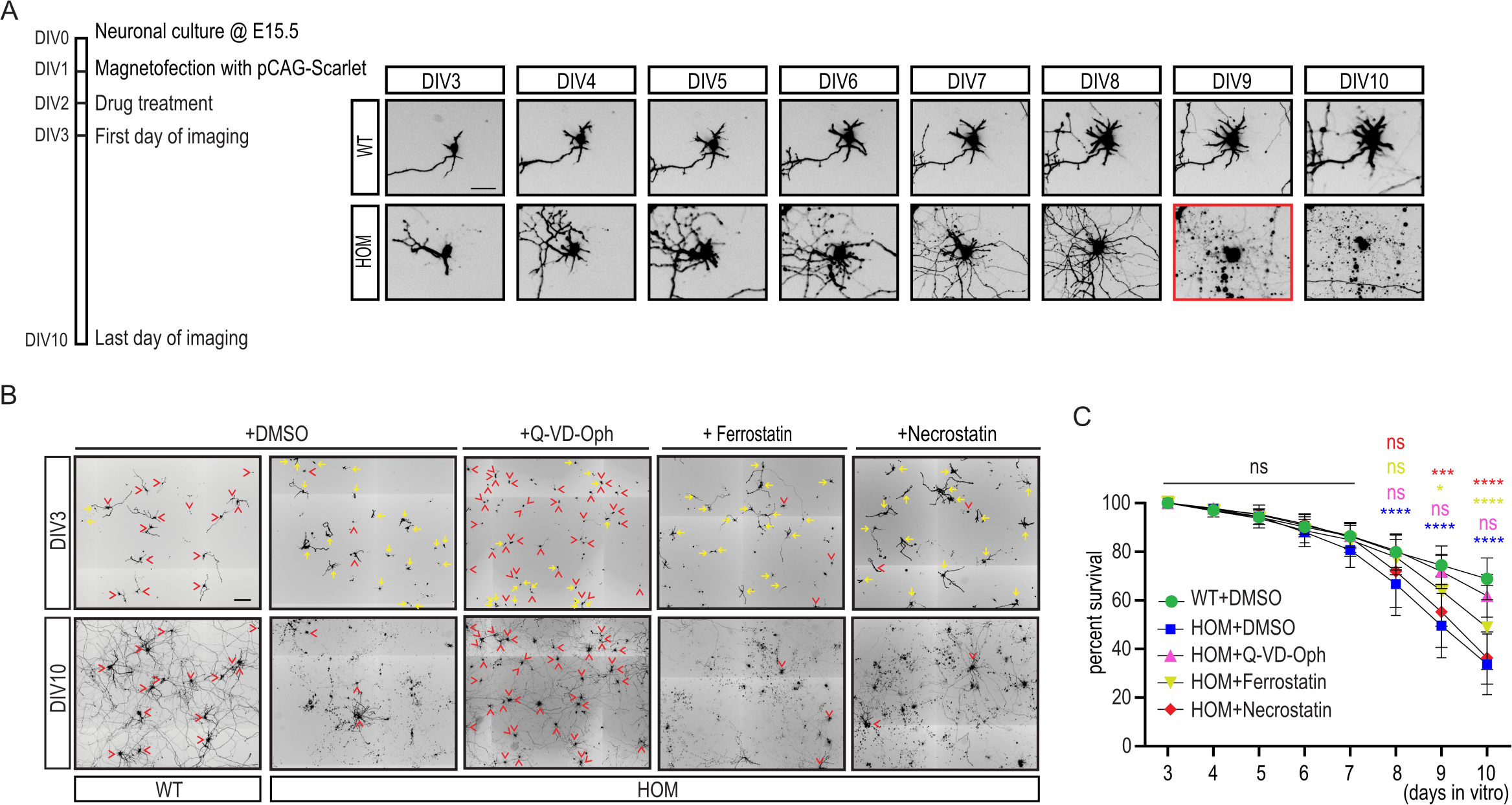
***Wdr47* deficient neurons die via apoptosis. (A)** *Left panel.* Schematic of the viability analysis in primary neuronal cultures. *Right panel.* Fluorescence imaging depicting single scarlet positive neurons (in black) that were followed from DIV3 to DIV10 in primary cultures obtained from WT and HOM (*Wdr47*^tm1b/tm1b^) embryos. Red rectangle indicates the time of neuronal death. Scale bar: 50 µm. **(B-C)** Neuronal survival *in vitro* in WT and HOM (*Wdr47*^tm1b/tm1b^) primary neuronal cultures upon treatment with different drugs. **(B)** Representative fields, at DIV3 and DIV10, of neuronal cultures treated with DMSO, Qvd-OPh (Caspase inhibitor), Ferrostatin (ferroptosis inhibitor) and Necrostatin (Necroptosis inhibitor) at DIV2. Scarlet positive electroporated neurons are depicted in black. Yellow arrows correspond to neurons that died, red arrowheads correspond to neurons that are alive and could be followed from DIV3 to DIV10. Scale bar: 200 µm **(C)** Survival of neurons from DIV3 to DIV10 upon different drug treatments. Data (means ± sd) from at least 3 independent cultures per condition was analyzed by two-way ANOVA, with Bonferroni’s multiple comparison test, ns, non-significant, *P < 0.05, ***P < 0.0005, ****P < 0.0001.

### *Wdr47*-deficient neurons present a neurodegenerative signature at the molecular level

To gain molecular insights into the mechanisms underlying neuronal death induced by the loss of *Wdr47,* we compared the global transcriptome of *Wdr47*^WT/WT^ (WT) and *Wdr47*^tm1b/tm1b^ knock-out (HOM) cortical neurons at DIV6, 2 days before the initiation of neuronal death using mRNA sequencing. Differential gene expression analysis identified 46 upregulated and 159 downregulated transcripts in HOM neurons (adjusted p value < 0.05) (**Figure 6A, Supplementary Table 2**). Gene Ontology (GO) enrichment analysis on differentially expressed genes (DEGs) revealed that, while upregulated genes were not enriched for any terms, downregulated genes were enriched in several biological process terms including regulation of synaptic plasticity (qvalue=0.005), regulation of neuron apoptotic process (qvalue=0.049), neuron projection extension (qvalue=0.042), postsynaptic specialization organization/assembly (qvalue= 0.0003) and regulation of microtubule polymerization or depolymerization (qvalue=0.047) (**Figure 6B**).

**Figure 6:**
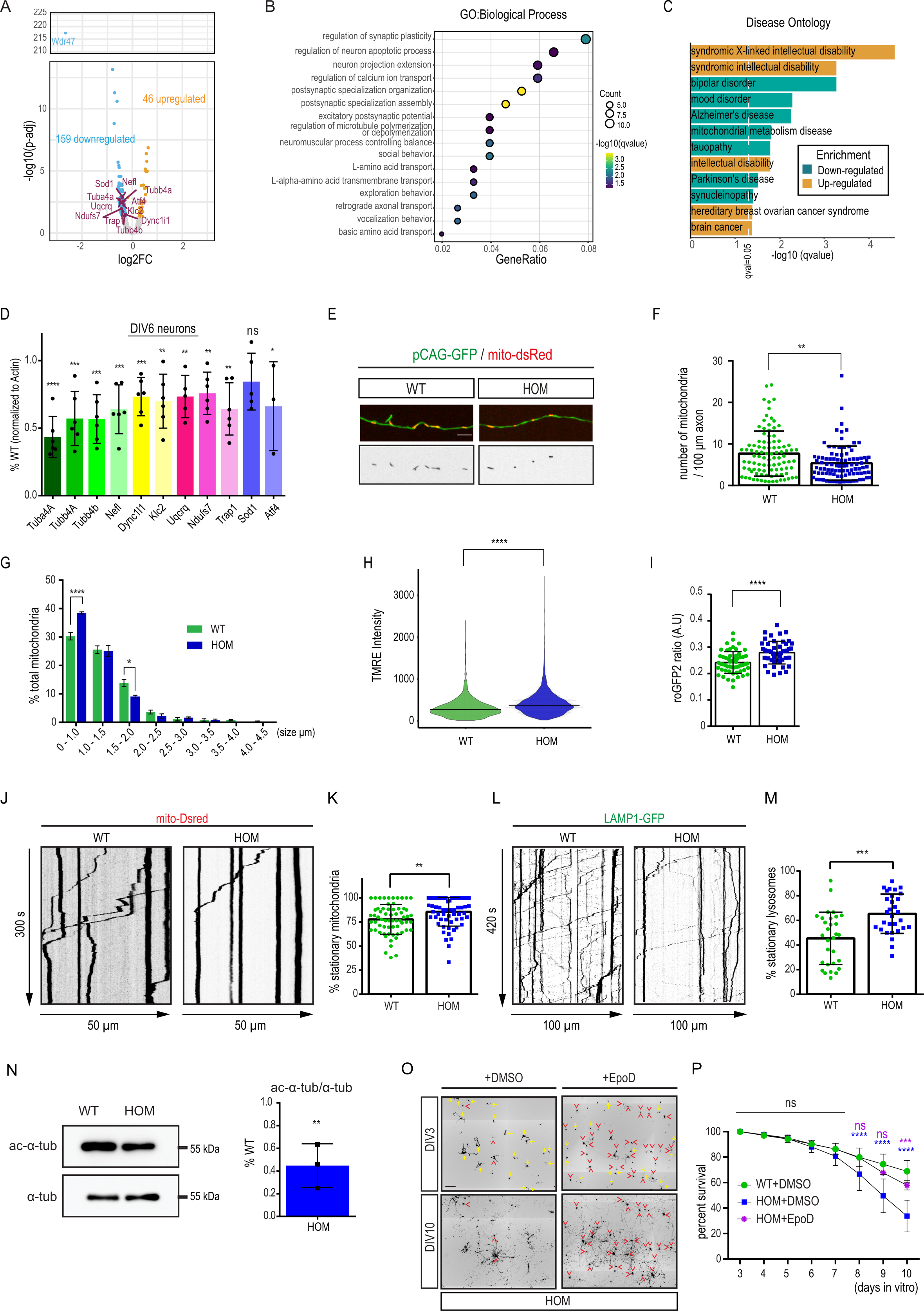
***Wdr47* deficient neurons present neurodegenerative hallmarks with altered mitochondrial and microtubule homeostasis. (A)** Volcano plot showing the negative log_10_ adjusted *P*value (p-adj) of all genes against their log_2_ fold change (log_2_FC) (*Wdr47*^tm1b/tm1b^ versus WT neurons). Up-regulated and down-regulated genes (p-adj < 0.05) are in orange and blue respectively and the neurodegeneration related genes selected for validation of RNA-seq results are labeled in red. **(B)** GO term analysis of downregulated genes in DIV6 neurons in *Wdr47*^tm1b/tm1b^ versus WT. The size of the circle represents the number of genes enriched in the GO term and the color of the circles represents the −log_10_ (qvalue). **(C)** Disease Ontology analysis of dysregulated genes in DIV6 neurons in *Wdr47*^tm1b/tm1b^ versus WT. Diseases enriched among upregulated and downregulated genes are depicted in yellow and blue respectively. Dashed line indicates qvalue equal to 0,05. **(D)** RT-qPCR validation of RNA-seq results for 11 downregulated genes associated to neurodegenerative diseases. Data (means ± sd) from at least 3 independent cultures per condition was analyzed by unpaired t-test. **(E-G)** Assessment of mitochondria in axons of WT and HOM (*Wdr47*^tm1b/tm1b^) DIV6 cortical neurons magnetofected with pCAG-GFP and mito-dsred. **(E)** Live-cell confocal imaging of individual mitochondria (red) in single axons (green) of WT and HOM (*Wdr47*^tm1b/tm1b^) DIV6 cortical neurons. Scale bar, 5 µm. **(F)** Number of mitochondria per 100 µm of axon in DIV6 cortical neurons (n >90 axons per condition). Data (means ± sd) from 5 independent cultures per condition was analyzed by unpaired t-test with Welch’s correction. **(G)** Quantitative assessment of mitochondrial morphology (Feret’s diameter) in axons of DIV6 cortical neurons (n > 400 mitochondria per condition). Data (means ± SEM) from 5 independent cultures per condition was analyzed by two-way ANOVA. **(H)** Mitochondrial membrane potential (ΔΨm) (TMRE intensity) of individual mitochondria in the neurites of WT and HOM (*Wdr47*^tm1b/tm1b^) DIV6 cortical neurons (n > 3500 mitochondria per condition). Data from at least 3 independent cultures was analyzed by Mann Whitney test. **(I)** Redox potential (mito-Grx1-roGFP2) of mitochondria in the neurites of WT and HOM (*Wdr47*^tm1b/tm1b^) DIV6 cortical neurons. (n >50 neurites per condition). Data (means ± sd) from 3 independent cultures per condition was analyzed by unpaired t-test with Welch correction. **(J-M)** Neurons were magnetofected at DIV4 with **(J-K)** mito-Dsred and **(L-M)** Lamp1-GFP to analyze mitochondria and lysosome motility respectively by videomicroscopy. Kymographs illustrate the motility of **(J)** mitochondria and **(L)** lysosomes in time (y, sec) and space (x, µm). Histograms represent the percentage of stationary **(K)** mitochondria (n>60 axons per condition from 5 independent cultures) and **(M)** lysosomes (n=30 axons per condition from 4 independent cultures). Data (means ± sd) was analyzed by unpaired t-test. **(N)** Western blot analysis shows decreased levels of acetylated tubulin in DIV6 HOM (*Wdr47*^tm1b/tm1b^) primary neurons compared to WT. Data (means ± sd) from 3 independent cultures was analyzed by unpaired t-test. **(O-P)** Neuronal survival in vitro in WT and HOM (*Wdr47*^tm1b/tm1b^) primary neuronal cultures upon treatment with MT stabilizer EpoD. **(O)** Representative fields, at DIV3 and DIV10, of neuronal cultures treated with DMSO and EpoD at DIV2. Scarlet positive electroporated neurons are depicted in black. Yellow arrows correspond to neurons that died, red arrowheads correspond to neurons that are alive and could be followed from DIV3 to DIV10. Scale bar: 200 µm. **(P)** Survival of neurons from DIV3 to DIV10 upon treatment with DMSO and EpoD (10nM). Data (means ± sd) from at least 3 independent cultures per condition was analyzed by two-way ANOVA, with Bonferroni’s multiple comparison test. ns, non- significant, *P < 0.05, **P<0.005, ***P < 0.0005, ****P < 0.0001.

Further enrichment analysis of disease ontology terms indicated on one hand that upregulated genes were mainly associated to neurodevelopmental deficits including various forms of intellectual disability and on the other hand that downregulated genes were linked to neurodegenerative diseases (*e.g.* Alzheimer’s and Parkinson’s disease) (**Figure 6C**), in line with a degenerative phenotype occurring at early stage.

In accordance, among the deregulated genes we found genes encoding for proteins involved in pathways critical to maintain neuronal homeostasis including intracellular trafficking (Cytoskeleton subunits, Tuba4A, Tubb4A, Tubb4B and Nefl; molecular motors, Dyncl1l and Klc2), mitochondrial function (Uqcrq, Ndufs7 and Trap1) and response to stress (Sod1 and Atf4) (**Figure 6A**). Downregulation of all those genes but Sod1 was confirmed by quantitative polymerase chain reaction (RT-qPCR) (**Figure 6D**) in *Wdr47*-deficient neurons at DIV6. Interestingly, in accordance with a sudden burst of cell death, none of those genes were differentially expressed in HOM cultures at DIV4 (**Supplementary** Figure 6A).

Collectively, these data highlighted several key cellular mechanisms whose combined deregulation upon loss of *Wdr47* likely contributes to rapid neuronal death.

### Loss of *Wdr47* impairs mitochondrial homeostasis in neurons

Given the presence of *Ndufs7* and *Uqcrq,* members of electron transport chain and *Trap1,* a mitochondrial chaperone that regulates mitochondrial homeostasis and bioenergetics among the downregulated genes in *Wdr47* KO neurons (**Figures 6A and 6D**) and the causative link between mitochondrial dysfunction and neuronal death ^23, 24^, we tested whether mitochondrial impairment could be one the mechanisms leading to the early death of *Wdr47*-deficient neurons. Live imaging of DIV6 WT and *Wdr47* KO neurons after transfection of pCAG-GFP and pCAG-mito-dsRed constructs to label axons and mitochondria respectively, revealed that *Wdr47* deficient axons contain fewer mitochondria (**Figures 6E- 6F**). In addition, the relative distribution of mitochondrial size is shifted towards smaller mitochondria in *Wdr47*-lacking neurons (**Figure 6G**), indicating an increased prevalence of fragmented mitochondria, a hallmark of damaged mitochondria. Then, to further assess mitochondrial integrity, we measured the mitochondrial inner membrane potential (ΔΨm) in WT and *Wdr47* KO neurons by performing dual imaging of TMRE (tetramethylrhodamine, ethyl ester) and mitotracker green, two dyes that allow labelling of mitochondria in a potential-dependent and independent manner respectively. Surprisingly, we found that in neurites of DIV6 *Wdr47*-deficient neurons, mitochondria showed 36% increased TMRE signal (mean TMRE intensity of 274.3 in WT and 373.4 in *Wdr47* deficient neurons) (**Figure 6H**) reflecting higher membrane potential. As mitochondrial hyperpolarization can lead to an increase in reactive oxygen species (ROS) production ^25^, we employed mitochondrial-targeted ratiometric roGFP2 probe (mito- roGFP2) ^26^ to measure the mitochondrial redox potential and observed a 15% increase of the roGFP2 ratio in *Wdr47*-deficient neurites compared to WT neurites indicating increased ROS production in mutant neurons (**Figure 6I**). Collectively, our results suggested that WDR47 regulates mitochondrial homeostasis, a function potentially required for its pro-survival activity.

### Compromised axonal transport and MT stability precede degeneration of *Wdr47*-deficient neurons

Given the critical role of axonal trafficking in neuronal homeostasis ^27^ and the deregulation of genes associated to axonal transport in *Wdr47*-KO neurons (**Figures 6A, 6B and 6D**), we hypothesized that Wdr47 survival-related function could also be mediated through regulation of axonal transport. We therefore analyzed axonal transport of mitochondria and lysosomes in WT and *Wdr47* deficient neurons after transfection of pCAG-mito-dsRed and Lamp1-GFP constructs respectively. Similar to previous reports ^28, 29^ in axons of WT neurons, 78% of mitochondria and 45% of lysosomes were stationary, whereas upon loss of *Wdr47*, the fraction of stationary mitochondria and lysosomes increased to 86% and 65 % respectively (**Figures 6J-6M**). Nonetheless, apart from a mild decrease in the velocity of anterograde moving mitochondria, velocity of the motile organelles was unchanged (**Supplementary** Figures 6B and 6C). As WDR47 has been shown to interact with and stabilize the microtubule (MT) cytoskeleton ^8, 9, 10^, we reasoned that the faulty axonal transport observed in *Wdr47*-deficient neurons (**Figures 6J-6M**) could stem from an unstable microtubule network. In line with this hypothesis, “*regulation of microtubule polymerization/depolymerization*” was among the deregulated GO biological processes (**Figure 6B**) and several tubulin members, the major component of MTs, were downregulated upon *Wdr47* deficiency in the RNA-seq and qPCR (**Figures 6A and 6D**). We therefore assessed the levels of stabilized (acetylated) and dynamic (tyrosinated) MTs in primary cultures of WT and *Wdr47*^tm1b/tm1b^ HOM neurons at DIV6 and showed that levels of acetylated tubulin were decreased by more than half (**Figure 6N**) while the total alpha tubulin and tyrosinated alpha tubulin levels were not altered (**Supplementary** Figure 6D), confirming the instability of the MT network upon *Wdr47* depletion. Finally, we tested the ability of Epothilone D (EpoD), a MT stabilizing compound ^30^, to rescue the survival defects induced by the loss of *Wdr47*. We treated WT and HOM neurons at DIV2 and monitored their survival over 10 days. We demonstrated that while it did not have any effect on the survival of WT neurons (**Supplementary** Figures 6E and 6F), EpoD treatment significantly recued the death of *Wdr47*- deficient neurons (34% and 58% survival in untreated and EpoD treated DIV10 HOM cultures respectively) (**Figures 6O and 6P**), indicating a major contribution of altered MT dynamics to the rapid degeneration of *Wdr47*-deficient neurons.

Altogether, these results indicated that WDR47 exerts its pro-survival function, at least in part, through the regulation of axonal transport and MT stability.

### *Wdr47* partial loss-of-function variants impede interhemispheric connectivity

Having validated the pathogenicity of the complete loss-of-function p.(Arg193His) variant using depletion of *Wdr47* in mice, and considering the presence of CCD in all patients, we hypothesized, that the other variants that do not directly impair WDR47 expression could lead to CC abnormalities through partial loss of function. To test this hypothesis, we tested the ability of p.(Asp466His), p.(Lys592Arg), p.(Pro650Leu) and p.(His659Pro) variants to restore CC anomalies induced by loss of *Wdr47* at P21 once callosal axons achieve their adult-like arborization pattern. We performed IUEs of NeuroD-Cre-GFP and pCAG2- mScarlet plasmids together with either WT or mutant WDR47 constructs in E15.5 *Wdr47*^fl/fl^ embryos and analyzed the scarlet-positive neurons and their axons. In contrast to the total loss of *Wdr47* that leads to an early onset burst of neuronal death (**Figure 3F**), all tested variants rescued the neuronal death phenotype as revealed by the bipolar morphology of electroporated neurons (**Figure 7A**) and the absence of pyknotic nuclei and Caspase 3 positive cells (data not shown). Nonetheless, expression of the p.(Pro650Leu) and p.(His659Pro) variants very slightly rescued axonal projection as shown by the reduced density of scarlet-positive axons in the midline compared to control (-71%, p=0.0003 and -67%, p=0.0011 respectively) (**Figures 7B-C**). Interestingly, the variants found as compound heterozygous in M03 have distinct effect after expression in *Wdr47*-depleted neurons. Indeed, while the expression of the p.(Lys592Arg) variant fully restore the CC thickness, the expression of the p.(Asp466His) variant partially rescued the density of scarlet positive axons in the midline (-45%, p=0.0133) (**Figures 7B-C**). Rerouting through alternate commissures was excluded as no aberrant axonal projections were observed and expression of none of the variants in the *Wdr47*^fl/WT^ animals used as controls caused any CC phenotype (**Supplementary** Figures 7A and 7B). Altogether these results indicated that while the variant that leads to a total loss of *WDR47* causes AgCC through an early burst of neuronal death, variants that lead to partial loss of protein function induces thinning of CC through other mechanisms acting possibly on axon generation or extension.

**Figure 7:**
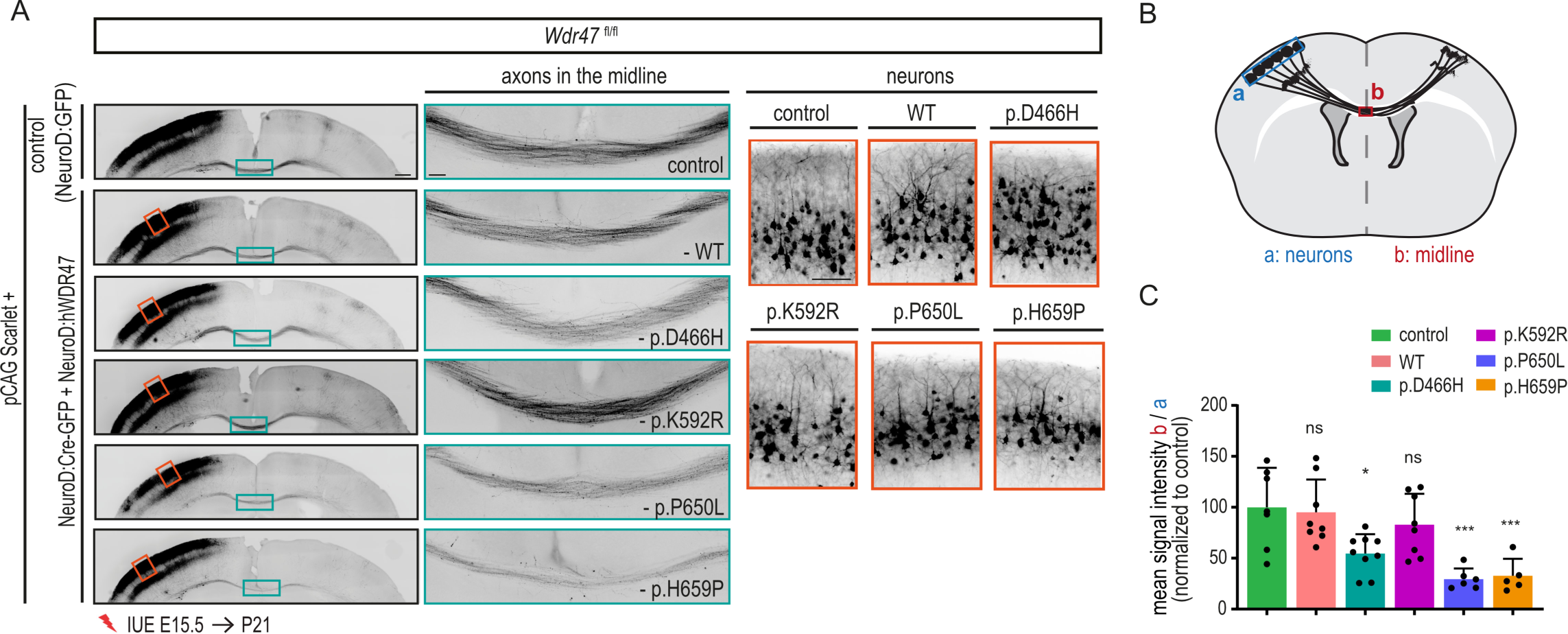
Different *WDR47* variants impede interhemispheric connectivity to varying degrees. (A) Coronal sections of P21 mouse brains electroporated at E15.5 with pCAG:Scarlet plasmid together with either a control plasmid or a NeuroD:Cre-GFP plasmid and a rescue construct (either a NeuroD: hWDR47-WT or a NeuroD:hWDR47 carrying one of the human mutations). Scarlet positive electroporated neurons are depicted in black. Close-up views of the green boxed area show that different constructs have different effects on rescue of CC phenotypes. Close-up views of the red boxed areas show bipolar morphology of the neurons in the cortical plate. Scale bars: 500µm, 100µm (green and red boxed insets). **(B)** Schematic describing the methods used to quantify **(C)** the CC thickness (mean intensity of the scarlet signal in the CC (red box) normalized on the mean intensity in the cortical plate (blue box)) **(C)** CC thickness upon introduction of different WDR47 mutated constructs. Data (means ± sd) from at least 5 pups from 2-3 different litters per condition were analyzed by one-way ANOVA, with Bonferroni’s multiple comparison test, ns, non-significant, *P < 0.05, ***P < 0.0005.

## DISCUSSION

In this study we demonstrate a causative link between human bi-allelic loss-of-function (LOF) missense variants in *WDR47* and neuroanatomical phenotypes (NAPs). In strong concordance with the previously characterized *Wdr47* deficient mouse models ^7^, corpus callosum dysgenesis (CCD) and microcephaly are fully penetrant features that are common to all human patients (5/5) (**Table 1**). Other non-fully penetrant NAPs include periventricular heterotopia (PVHT), a neuronal migration disorder, that reflects the regulatory function of Wdr47 in migration ^7, 9^ and hydrocephalus, that has been observed in mouse models invalidated for *Wdr47* specifically in precursors of ependymal cells ^16^.

We previously showed, using constitutive knock-out models, that *Wdr47* deficiency leads to reduced number of progenitors and increased neuronal death both of which likely contribute to the microcephaly phenotype ^7^. Here, we addressed the cellular and molecular origins of CCD. Development of CC requires the interplay of different cellular populations including midline glial structures, meninges, pioneering axons, cortical progenitors and callosal projection neurons (CPNs) ^5^. Using specific inactivation of *Wdr47* in neurons, we identified a major contribution of post mitotic neurons to the CC phenotype (**Figures 2 and 3 and Table 2**). Notably, upon acute deletion of *Wdr47* in CPNs, in addition to the axon extension defects at P2, we observed a sudden massive induction of cell death at P3, with nearly no neurons remaining and CC being totally lost at P8. Such a robust burst of neuronal death was not anticipated based on previous studies that rather indicate that human CCD syndromes are caused by faulty neuronal specification and migration, axon extension and guidance ^5^.

Our study identified five missense variants that are distributed throughout the protein. The Arginine residue at position 193, found to be mutated in M01 and M02, is located within the Wdr47 N-terminal domain (WDR47-NTD), in the cross over region (COR), a subdomain involved in the formation of the WDR47-NTD intertwined dimer, that serves as an interaction interface with protein partners ^14^. Thanks to the crystal structure of the mouse WDR47-NTD ^14^, we showed that the Arginine 193 residue forms a salt bridge with the glutamate 205 located in the COR of the other protomer (**Figure 1J**). Notably, the substitution of the Arginine 193 with a Histidine disrupts the salt bridge between the two protomers and likely destabilizes the dimer. In accordance with this hypothesis, we showed that the p.(Arg193His) variant leads to a dramatic reduction of WDR47 expression at the protein level specifically (**Figures 1K- M**). In line with the phenotype of the constitutive *Wdr47* KO mice ^7, 9^, the two patients, M01 and M02, present with the severest phenotype with neonatal death occurring 24-48 hours after birth and agenesis of the corpus callosum. Given the NTD-mediated formation of the WDR47 intertwined dimer, it is conceivable that mutations in this domain that interfere with the formation of WDR47 dimer could be highly deleterious for WDR47 expression and/or function and lead to lethality *in utero* or shortly after birth. Supporting an intolerance to complete loss of function, no homozygous *WDR47* frameshift variants have been reported in the gnomAD (Genome Aggregation Database, v4.0.0) collection of populations.

Interestingly, we also reported patients with variants located outside the NTD presenting with milder CC phenotypes. Using acute depletion of *Wdr47* in neurons and complementation assays, we showed that p.(Pro650Leu), p.(His659Pro) and p.(Asp466His) variants lead to CC abnormalities not by primarily affecting the neuronal survival but possibly by impairing axon generation and growth (**Figures 7A-C)**.

Additionally, our results demonstrate that *Wdr47* haploinsufficiency does not lead to any survival or axon extension defects (**Figures 3A-F and Supplementary** Figures 3A-B). Altogether these results led us to propose a model in which a minimal level of WDR47 activity is required to ensure proper development of the corpus callosum in human and mice. In this model, maintaining 50% of WDR47 function is sufficient to promote CC development. Below this threshold, the more functions are lost, the more severe and broad are the phenotypes. Consistently, total loss of function (p.(Arg193His)) induces an early lethality and a complete loss of CC, while partial loss of function (p.(Pro650Leu), p.(His659Pro) and p.(Asp466His), seen as either a decrease of all known activities or a loss of only some specific function, leads to thinning of the CC likely through defects in axon extension.

At the cellular level, we demonstrated that the pro-survival function of WDR47 is independent of its roles in neuronal migration and axon extension. Indeed, although the introduction of Camsap3 restored the migration and axon extension deficits as previously suggested ^9^, it failed to rescue the neuronal survival defects (**Figure 4**). These results also indicate that: 1) an early onset burst of neuronal apoptosis is the main driver for the CC phenotype upon *Wdr47* loss and 2) WDR47 requires different downstream effectors other than Camsap3 to mediate neuronal survival function of WDR47. Given the presence of WDR47 in subcellular fraction enriched in synaptic vesicles ^31^ and the perturbations of several genes related to synapse function/organization in *Wdr47* deficient neurons (**Figure 6B**), one can anticipate a possible synaptic function of WDR47. Nevertheless, as *Wdr47* deficient neurons die at P3, when synaptic maturation is not fully established, it is unlikely that a synaptic dysfunction is triggering the loss of *Wdr47* deficient neurons *in vivo*.

Our study further provides insights into the mechanisms underlying the rapid degeneration of *Wdr47* deficient neurons. Several evidence point towards a role of WDR47 on the regulation of microtubule (MT) homeostasis and intracellular organelles trafficking. First, *Wdr47* deficient neurons had remarkably reduced acetylated tubulin, indicative of an unstable MT network ^32^ (**Figures 6N-P**). The possible role of WDR47 in stabilizing microtubule might be mediated through its binding to two known regulators of MT dynamics: i) the superior cervical ganglion-10 (SCG10) protein, a well-established microtubule-destabilizing protein ^7, 33^; and/or ii) the MT-associated protein 8 (MAP8) that stabilizes MT and serves as an intermediate to bring WDR47 to the MTs ^8, 34, 35^. Although Camsap3, known to interact with Wdr47 and stabilize MT minus ends ^9, 10^, could mediate Wdr47 MT dynamic regulatory function, it unlikely contributes to the pro-survival function of WDR47, as it failed to rescue callosal neurons survival *in vivo* (**Figure 4D**). Second, we reported impaired MT-based transport of lysosomes and mitochondria in *Wdr47* deficient neurons (**Figures 6J-M and Supplementary** Figure 6B). Although axonal transport defects of *Wdr47* deficient neurons can be at least partially attributed to decreased acetylation and defects in MT stability ^32, 36^ (**Figures 6N-P**), unpublished co-immunoprecipitation assays showed that WDR47 can interact with motor proteins dynein and kinesin and hence possibly influence interaction of cargos with motor proteins. Accordingly, WDR47 protein has been found at the surface of motile vesicles isolated from mouse cerebral cortex ^37^. Moreover, recent studies showed that WDR47 negatively controls the formation of motile molecular motor complexes, thereby regulating MT-dependent transport of ATG9A containing vesicles ^11^ and intraflagellar transport of cargoes along the ciliary axoneme ^38^. Finally, we demonstrated that treatment of *Wdr4*7 deficient neurons with EpoD, a MT stabilizing compound ^30^ that favors intracellular transport ^39^ significantly improved the survival of *Wdr47* deficient neurons (**Figures 6O,P**).

Whereas one cannot entirely exclude that lack of full rescue with EpoD is due to dose and/or time dependent reasons, other underlying mechanisms besides decreased MT stability, can also contribute to the rapid degeneration of *Wdr47* deficient neurons.

In accordance, we unveiled a previously unidentified function of WDR47 in maintenance of mitochondrial homeostasis. We showed that loss of *Wdr47* leads to fragmentation of mitochondria and increased production of reactive oxygen species (ROS) (**Figures 6E-I**). Although these processes are usually associated to depolarization of mitochondria ^40, 41^, we unexpectedly observed a hyperpolarization in *Wdr47*-deficient neurons (**Figure 6H**). Interestingly, such hyperpolarization of mitochondrial membrane has been reported to lead to oxidative stress and neuronal death in patient derived neurons with mutations in Tau ^25^. Given that WDR47 interacts with mitochondrial proteins in neurons ^42^, it is tempting to speculate that, in addition to its function in MT-related transport of mitochondria, WDR47 could have a direct function on mitochondrial homeostasis. Increased ROS and decreased levels of *Sod1,* a critical antioxidant enzyme, in *Wdr47* deficient neurons reflects an imbalance between pro-oxidant species and antioxidant systems (**Figures 6D, I**). As neurons are highly vulnerable to oxidative stress owing to their non-dividing nature and high metabolic demand ^43^, loss of redox homeostasis could be an additional potential trigger for the loss of *Wdr47* deficient neurons. However, our attempts to rescue the survival defects by reducing ROS in *Wdr47* deficient neurons using an antioxidant N-acetyl cysteine (NAC) failed (data not shown). Yet, excessive ROS levels might in turn result in mitochondrial DNA/proteins damages, inner mitochondrial membrane permeabilization and release of cytochrome c into the cytosol that all ultimately activate the apoptotic machinery ^44, 45^ and that might not be rescued by ROS scavenging. In addition, previous studies that use a plethora of antioxidants to target oxidative stress in different models of neurodegeneration suggests that the efficiency of different antioxidants varies across models and that a combination of different antioxidants better attenuate the damaging effect of ROS ^44, 45^. Therefore, before one can firmly conclude that neuronal death is not triggered by oxidative stress, additional rescue strategies with different classes antioxidants alone or in combination should be tested.

Whether WDR47 variants in different domains would interfere with specific functions to different extends remained to be investigated. MT-stability function could rely on both N- and C-terminal domains as the WD40 domain has been previously involved in the binding of WDR47 to MT via MAP8 ^8^, while the N-terminal domain is indispensable for binding to SCG10 ^7^. Given the strong homology to LIS1, that presents a N-terminal dimerization domain and a WD40 C-terminal domain required for its binding to molecular motors ^46, 47^, WDR47 C-terminal domain appears as a strong candidate to mediate its function in regulation of intracellular transport. However, use of sensitive methods to precisely map the function of WDR47 is required to unveil possible pleiotropic roles of the different WDR47 domains and potentially better predict the pathogenicity of variants depending on the position within the WDR47 protein sequence.

Notably, the molecular alterations that we reported in *Wdr47*-deficient developing neurons are reminiscent of pathological mechanisms delineating the ultimate loss of neurons in neurodegenerative diseases ^23, 24, 27, 48, 49^. We report perturbations of a significant number of Alzheimer’s and Parkinson’s related genes in *Wdr47* deficient primary neuronal cultures, corroborating a recent study that identified *WDR47* as a hub gene in Alzheimer’s ^12^. In addition, loss of MT homeostasis, impaired transport of cargoes and mitochondria alteration are hallmarks of a variety of common neurodegenerative diseases, such as Alzheimer’s disease (AD), amyotrophic lateral sclerosis (ALS), Parkinson’s disease (PD) and Huntington’s disease. Very interestingly, Wdr47 shares similarities with Huntingtin (Htt) and Tau, the proteins whose dysfunction cause HD and AD/frontotemporal dementia (FTD) respectively. First, alike Htt ^50, 51^ and Tau ^52, 53^, there is growing evidence for a role of WDR47 in regulation of mitochondria homeostasis, in addition to its initially recognized MT-associated function. Second, similar to the ability of Htt and Tau to interact with mitochondrial proteins ^54, 55, 56^, two recent analysis of the WDR47 interactome revealed binding of WDR47 to mitochondrial proteins involved in ATP synthase complex, pyruvate metabolism and transport of metabolites and ions ^9, 10^. Therefore, LOF mutations in *Wdr47* could impair mitochondrial bioenergetics by disrupting its interaction with mitochondrial proteins and contribute to an early neurodegenerative phenotype. Third, WDR47 was identified as a binding partner of Huntingtin ^57^.

Given the growing evidence that neurodegeneration in Huntington’s disease stem from alterations occurring during development ^58, 59, 60^, it is tempting to speculate that WDR47 safeguards callosal neurons from early activation of neurodegenerative mechanisms.

In summary, our data identify *WDR47* as an important gene for neuroanatomical disorders including CCD and microcephaly in humans and mice. We show that degeneration of *Wdr47*-deficient neurons is the main driver of CC agenesis upon *Wdr47* loss and WDR47 safeguards callosal neurons from early-onset neuronal death by contributing to the maintenance of mitochondrial and microtubule homeostasis. We suggest that screening of *WDR47* should be considered in unexplained cases of corpus callosum abnormalities and microcephaly.

## MATERIALS AND METHODS

### Whole exome sequencing (WES)

#### Patients M01 and M02

Research human subjects were enrolled in an IRB-approved research protocol with informed consent. Blood was collected from index and available relatives in EDTA tubes for DNA extraction. All subjects were exome sequenced and the resulting variants were filtered by the candidate autozygome as described previously ^61, 62^.

#### Patients M03

DNA was extracted from peripheral blood of the patients and their parents. Patient M03’s DNA was captured with the SureSelect Human All Exon V6 Kit (Agilent Technologies, Santa Clara, CA, USA) and sequenced on a NextSeq500 (Illumina, San Diego, CA, USA) with 150-bp paired-end reads. Reads were aligned to reference genome (GRCh38) using BWA-MEM (Version 0.7.17). Duplicated reads were removed by Picard (Version 2.20.3), and local realignment and base quality recalibration were performed by GATK (Version 4.1.9.0). The total aligned bases and mean depth of coverage against RefSeq coding sequences (33.9 Mb) were 2,278,284,739 bp and 67.18. The target RefSeq coding sequences covered by greater than 20 reads were 95%. Variants were identified with the GATK HaplotypeCaller. The final variants were annotated with Annovar (Wang et al. 2010) for predictive value of functional impact of the coding variants and assessing allele frequency using follow databases: the gnomAD database v3.1.2 (https://gnomad.broadinstitute.org/), the Human genetic variation database (http://www.hgvd.genome.med.kyoto-u.ac.jp/), and the ToMMo 8.3KJPN Allele Frequency Panel (v20200831) (https://jmorp.megabank.tohoku.ac.jp/202102/downloads/legacy/). We focused on rare variants with minor allele frequencies below 1% in the above four databases. The damaging prediction was performed by SIFT, Polyphen-2, MutationTaster, CADD and M-CAP programs. Candidate variants were confirmed by Sanger sequencing using an ABI 3130xl Genetic Analyzer (Applied Biosystems, Foster City, CA).

#### Patient M04

Genomic DNA was fragmented, and the exons of the known genes in the human genome, as well as the corresponding exon-intron boundaries were enriched using Roche KAPA capture technology (Basel, Switzerland), amplified and sequenced simultaneously by Illumina technology (San Diego, US). The target regions were sequenced with an average coverage of 148.9-fold. For about 99.9% of the regions of interest a 15-fold coverage was obtained. NGS data were aligned to the hg19 genome assembly. Variant calling and annotation were performed by an in-house developed bioinformatics pipeline. Identified SNVs and indels were filtered against external and internal databases focusing on rare variants with a minor allele frequency (MAF) in gnomAD of 1% or less and removing known artifacts and variants in regions with highly homologous regions. The filtering of the exome data targeted X-linked, autosomal dominant and recessive disorders. In parallel, we received parental samples for which WES was performed in a Trio setting. Classification of variants was conducted based on ACMG guidelines ^63^ considering variant databases including but not limited to ClinVar, bioinformatics prediction tools and literature status. A change of pathogenicity classifications over time cannot be excluded. Variants annotated as common polymorphisms in databases or literature or that were classified as (likely) benign were neglected.

#### Patient M05

WES was performed at Mendelics Genomic Analysis facilities using Illumina NovaSeq 6000. Sequencing library was built with Illumina Nextera Flex and for capture of target regions Customized Exome Kit from Twist Biosciences is used. Sequencing of sample results in paired 101 bp sequences that are mapped to hg38 reference using BWA MEM software (http://bio-bwa.sourceforge.net/). 99.8% of the targeted regions had at least 10 reads sequenced, resulting in a total 43,754,430 generated and with a medium vertical coverage of 79-fold. Resulting BAM files were genotyped using Broad Institute best practices with GATK (https://software.broadinstitute.org/gatk/). Resulting VCF files were processed using Mendelics in-house pipeline for annotation and filtering. *In silico* pathogenicity evaluation was performed using VSA, a Mendelics proprietary machine-learning based software. Aligned BAM files were also processed by ExomeDepth (an R package, see https://cran.r-project.org/web/packages/ExomeDepth/index.html) in order to identify CNVs. Generated data was analyzed by Mendelics medical team and guided by clinical information provided by the primary physician. For analysis, several genomic tools and variant/phenotype databases were applied, including University of Santa Cruz Genomic Browser (https://genome.ucsc.edu/index.html), gnomAD browser v2.1.1 and v3.1.2 (https://gnomad.broadinstitute.org/), Human Gene Mutation Database – HGMD (http://www.hgmd.cf.ac.uk/ac/index.php), ClinVar (https://www.ncbi.nlm.nih.gov/clinvar/), Mastermind by Genomenom (https://mastermind.genomenon.com/); Leiden Open Variation Database – LOVD (https://www.lovd.nl/), OMIM – Online Mendelian Inheritance in Man (https://www.omim.org/), as well as medical literature by searching Pubmed (https://pubmed.ncbi.nlm.nih.gov/). Multiple parameters were considered during variant analysis - population frequency, gene associated phenotypes and inheritance pattern, presence of previous reports in medical literature and variant databases, functional data, segregation analysis and *in silico* predictions. Potentially clinically significant variants were visualized using Integrative Genomics Viewer - IGV Browser (https://software.broadinstitute.org/software/igv/).

Whenever parental consanguinity was present and no clinically relevant variants were found in genes with a presently known phenotype, search for rare homozygous variants in genes without an associated phenotype was performed, using MAF<= 1% as a filtering parameter. Genes in which potentially relevant variants were identified, were evaluated for expression pattern, biological function (when known), functional data and animal models that might be reminiscent of the patient’s phenotype, when available. Based on these parameters, the *WDR47* missense variant was considered a possible candidate and thus was validated in the proband and confirmed in the heterozygous state in both of his parents by Sanger sequencing.

### Mice

All animal studies were conducted in accordance with French regulations (EU Directive 86/609 – French Act Rural Code R 214-87 to 126) and all procedures were approved by the local ethics committee and the Research Ministry (APAFIS#15691-201806271458609 for *in utero* electroporation, APAFIS #15169 – 2018052111497498 for intracardiac final perfusion, APAFIS #2016010717527861 for neuroanatomical analysis). Mice were bred at the animal facility of the Mouse Clinical Institute (MCI) (IGBMC) or of the INSERM unit 1231 under controlled light/dark cycles and were provided with food (chow diet) and water ad libitum, with cage enrichment.

*Wdr47^tma(EUCOMM)Wtsi^*and *Camsap3^tma(EUCOMM)Wtsi^*knock-out mice generated by homologous recombination in embryonic stem cells using the knock-out-first allele method ^64^ **(Supplementary** Figure 2A) were obtained through collaboration with the Mouse Genetics Project (MGP) from the Welcome Sanger Institute (Cambridge, UK), a partner of the International Mouse Phenotyping Consortium (IMPC).

To obtain timed-pregnant females: hybrid F1 females *Wdr47*^tm1c/wt^ ^(fl/wt)^ were obtained by mating inbred 129S2/SvPasOrlRj males (Janvier-labs) with *Wdr47*^tm1c/tm1c(fl/fl)^ females (C57BL/6NCRL background Charles River Laboratories). F1 females were then crossed with *Wdr47*^tm1c/tm1c(fl/fl)^ males for *in utero* electroporation with dating of pregnancy by vaginal plug.

Nex^Cre^ mice ^17^ were obtained from Charles River and Nex^Cre^ mice were crossed with *Wdr47*^fl/fl^ mice to generate double mutant mice Nex^Cre/wt^*;Wdr47^fl/wt^* and those double mutant mice were crossed to generate mice with a deletion of *Wdr47* in post-mitotic neurons (Nex^Cre/wt^*;Wdr47*^fl/fl^ and Nex^Cre/Cre^*;Wdr47*^fl/fl^ and their controls (Nex^Cre/wt^*;Wdr47*^WT/WT^ and Nex^Cre/Cre^*;Wdr47*^WT/WT^).

CaMKIIα^Cre^ mice ^65^ were obtained from the Jacksons Laboratory and crossed with *Wdr47^fl/fl^*mice to generate double mutant mice CaMKIIα^Cre^; *Wdr47*^fl/WT^. Those double mutant mice were then crossed to generate mice with deletion of *Wdr47* in peri-natally born cortical neurons CaMKIIα^Cre^; *Wdr47*^fl/fl^ and their controls (CaMKIIα^Cre^*;Wdr47*^WT/WT^).

### Genotyping and PCR

Genotyping was done as follows: Genomic DNA was extracted from tail biopsies using PCR reagent (Viagen) supplemented with Proteinase K (1 mg/mL), heated at 55°C for 2-5 h. Proteinase K was inactivated for 45 min at 85°C, and cell debris was removed by centrifugation. PCRs for each line were carried as follows and presence of expected products were checked on 1% agarose gel.

PCR for each line was done as follows:

*Wdr47*^tm1a/WT^: *Wdr47* forward: 5’-TCCTTTGCTAACTTCCACTATCC-3’ and *Wdr47* reverse: 5’- TCAGCCTGGTCTACAGAGTTA-3’ for *Wdr47* targeted exon amplification. The presence of the wild type and knock-out alleles was indicated by 615 bp and 533 bp products, respectively.

*Wdr47*^tm1b/WT^: *Wdr47* forward: 5’-AGGTTGTCATGCAGTCTGGG-3’ and *Wdr47* reverse: 5’- GGATGACTATAAAGCGGTGCAAG-3’ for *Wdr47* targeted exon amplification; 5’- CTCCTACATAGTTGGCAGTGTTTGGG-3’ for knock-out allele. The presence of the wild type and knock- out alleles was indicated by 642 bp and 362 bp products, respectively.

*Wdr47*^tm1c/WT^: *Wdr47* forward: 5’-AGGTTGTCATGCAGTCTGGG-3’, *Wdr47* reverse: 5’- GGATGACTATAAAGCGGTGCAAG-3’ for *Wdr47* targeted exon amplification. The presence of the wild type and knock-out alleles was indicated by 642 bp and 848 bp products, respectively.

*Camsap3* ^tm1a/WT^: PCR genotyping was carried out using genomic DNA isolated from mouse tail samples by identifying the inserted cassettes, as described in ^66^.

Nex^Cre^: Nex^Cre^ forward: 5’-GAGTCCTGGAATCAGTCTTTTTC-3’, Nex cre WT reverse: 5’- AGAATGTGGAGTAGGGTGAC-3’, Nex^Cre^ KI reverse: 5’-CCGCATACCAGTGAAACAG-3’, for Nex targeted exon amplification. The presence of the wild type and knock-in alleles was indicated by 770 bp and 525 bp products, respectively.

CaMKIIα^Cre^: Cre forward 5’-ATCCGAAAAGAAAACGTTGA-3’, Cre reverse 5’- ATCCAGGTTACGGATATAGT -3’. Presence of the cre transgene was detected with a 500bp product.

### Neuroanatomical characterization of adult murine brains

To obtain the brain samples derived from *Wdr47*^tma(EUCOMM)Wtsi^ and *Camsap3*^tma(EUCOMM)Wtsi^ knock-out mice, adult mice (16 weeks old) were anesthetized using either Ketamine (100 mg/kg, intraperitoneally) and Xylazine (10 mg/kg, i.p.) and then the brains were dissected out and drop fixed in 10% neutral buffered formalin. Neuroanatomical studies were carried out using 3 homozygous *Camsap3*^-/-^ and 24 matched baseline WT mice as well as 3 homozygous *Wdr47*^tm1a/tm1a^ and 3 littermate WT mice, all on a C57BL/6NTac pure genetic background at 16-weeks of age. To facilitate data integration, CaMKIIα^Cre^; *Wdr47^fl/fl^* conditional mice were studied at 16 weeks of age on the same genetic background. We calculated statistical power using a recognized model (Gpower) validated for comparison of small numbers of mice (n=3) to evaluate neuroanatomical phenotypes with an effect size of 10% or more with a detection power of 80% ^4^. Paraffin embedded brain samples were cut at 5μm thickness using a sliding microtome (Micom HM 450) to obtain brain sections at Bregma -1.34 mm (*Camsap3*) or at Lateral +0.72 mm (*Wdr47*), according to the Allen Mouse Brain Atlas ^67^. Sections were stained with 0.1% Luxol Fast Blue (Solvent Blue 38; Sigma-Aldrich) and 0.1% Cresyl violet acetate (Sigma-Aldrich) and scanned using the high-resolution digital slide scanner (Nanozoomer 2.0HT, C9600 series) at 20× resolution. For *Camsap3*, three measurements were taken on the coronal plane and included the soma of the corpus callosum connecting the lateral part (lat) and the medial part (med) of the cerebral cortex, as well as the area of the dorsal hippocampal commissure (dhc). For *Wdr47*, we used a previously described procedure involving 40 measurements ^21^.

### Neuroanatomical characterization of embryonic murine brains

For male and female embryos at E18.5 (n=5 WT vs n=4 *Wdr47*^tm1a/tm1a^; n=6 WT vs n=6 Nex^Cre^; *Wdr47*^fl/fl^), the animals were kept cold on ice and killed by making a cut between the brain and spinal cord junction, before fixation in Bouin solution for 48 hours. The embryonic brains were then harvested, transferred to 70% ethanol, and manually embedded in paraffin using the following steps: three incubation baths in 70% ethanol for 30 minutes each, two baths in 95% ethanol for 30 minutes each, two baths in 100% ethanol for 45 minutes each, three baths in Histosol Plus for 1 hour each, and five baths in warm paraffin (60°C) for 30 minutes each, followed by incubation in warm paraffin overnight before casting in a mold. Brains were cut at a thickness of 5μm on a microtome (HM 450, Microm Microtech, France), such that we obtain sections matching planes at precise embryonic coronal sections, as explained in ^18^. The sections were stained with 0.1% Cresyl violet acetate (Sigma-Aldrich) and scanned using Nanozommer 2.0HT, C9600 series at 20× resolution.

Each image was quality controlled to assess whether (i) the section is at the correct position, (ii) the section is symmetrical, (iii) the staining is of good quality, and (iv) the image is good quality. Only images that fulfilled all of the quality control checks were fully processed. These quality control steps are essential for the detection of small to moderate neuroanatomical phenotypes and without which the large majority of neuro-anatomical phenotypes would be missed. This is explained in great details elsewhere ^18^. 54 brain morphological parameters made of area and length measurements were taken for each sample at E18.5 (**Supplementary Table1**). Assessed embryonic brain regions included the total brain area, the cortices (motor, insular, somatosensory, retrosplenial granular and motor), the hippocampus, the genu of the corpus callosum, the internal capsule, the caudate putamen, the fimbria of the hippocampus, the anterior commissure and the ventricles (lateral and third). Every aspect of the procedure was managed through a relational database using the FileMaker (FM) Pro database management system (detailed elsewhere ^68^). A list of histological parameters is also provided in **Figure 2**. Data were analyzed using two-tailed Student’s *t*-tests of equal variances.

### Plasmids and cloning

Wild-type (WT) human *WDR47* cDNA corresponding to UniprotKB O94967-3 isoform of WDR47 was obtained from DNASU and subcloned by restriction-ligation into the NeuroD-IresGFP ^69^ and the pcDNA3.1+/C-HA vectors. Constructs carrying the human *WDR47* variants p.(Arg193His), p.(Asp466His), p.(Lys592Arg), p.(Pro650Leu) and p.(His659Pro) were created from the WT CDS by Sequence and Ligation Independent Cloning (SLIC). Truncated constructs were generated from the NeuroD-WT- hWDR47-IresGFP and the pcDNA3.1-hWDR47-C-HA vectors by deletion of the LISH (amino acids 10- 42), CTLH (amino acid 45-102) and the WD40 (amino acids 658-920) domains by SLIC. pFASBac+GFP- Camsap3 plasmid was obtained from Addgene. GFP fused to the N-terminal of Camsap3 was removed and subcloned by restriction-ligation into the NeuroD-Ires-GFP vector. Neuro-D:Cre-GFP was a gift from L. Nguyen (GIGA, Liege, Belgium) and pCAG2-mScarlet vector was provided by J. Courchet (INMG, Lyon, France). Plasmid DNAs used in this study were prepared using the EndoFree plasmid purification kit (Macherey Nagel).

### *In utero* electroporation

The uterine horns of time pregnant (E14.5 and E15.5) mice were exposed after anesthetizing the females with isoflurane (2 L per min of oxygen, 4% isoflurane in the induction phase and 2% isoflurane during surgery operation; Tem Sega) and one ventricle of each embryo was injected with 0.5-1 μL of Endofree plasmid DNA solution mixed with 0.05% Fast Green (Sigma Aldrich) using pulled glass capillaries (Harvard apparatus, 1.0OD*0.58ID*100mmL) and a micro injector (Eppendorf Femto Jet). For migration analysis 3μg/μL of NeuroD:Cre-GFP vector together with 1μg/μL of either empty NeuroD-IRES-GFP or a rescue construct under the NeuroD promoter was used at E14.5. For CC analysis, the same plasmid mixes together with 0,8 µg/µl of pCAG2-mScarlet were used at E15.5. Plasmids were then further electroporated into the neuronal progenitors adjacent to the ventricle by discharging five electrical pulses of 40V and 50ms at 950 ms intervals using 3mm platinium tweezers electrodes (Sonidel CUY650P3) and ECM-830 BTX square wave electroporator (VWR international). After electroporation, embryos were placed back in the abdominal cavity and the abdomen was sutured using surgical needle and thread.

### Mouse brain fixation and cutting

For migration and CC analysis after IUE, E18.5 to P8 animals were sacrificed by head sectioning and P21 pups were sacrificed by terminal perfusion by 0.09% NaCl followed by 4% PFA. All brains were then fixed in 4% paraformaldehyde (PFA, Electron Microscopy Sciences) in Phosphate buffered saline (PBS, HyClone) overnight at 4°C. After fixation, brains were rinsed and embedded in a solution of 4% low-melting agarose (Bio-Rad) and cut into coronal sections (60 µm-thick for E18.5 and P2 mice, 100 μm- thick for P3 to P21 mice) using a vibrating-blade microtome (Leica VT1000S, Leica Microsystems).

Sections were maintained in PBS-azide 0,05% for short-term storage and in an Antifreeze solution (30% Ethyleneglycol, 20% Glycerol, 30% DH2O, 20% PO4 Buffer) for long-term storage.

### Immunolabeling

Vibratome sections were rinsed in PBS, permeabilized and blocked in 5% Normal Donkey Serum/PBST (1X PBS, 0.3% Triton X-100) and incubated with primary antibodies (**Supplementary Table 3**) diluted in blocking solution at 4°C overnight. After washing with PBST, the sections were incubated for 1h hour at room temperature with fluorescence-conjugated secondary antibodies coupled to Alexa-488 or Alexa-647 (Thermo Fischer Scientific). Nuclei were counterstained with Dapi (1µg/mL Sigma-Aldrich) and sections were mounted using Aquapolymount mounting solution (Polysciences Inc.). The slides were stored in the dark at 4°C. The primary and secondary antibodies used are listed in **Supplementary Table 3**.

### Plasmids, strains, media, and methods for yeast cells

The human *WDR47* cDNA either full-length (FL), or lacking the CTLH (ΔCTLH) and LISH (ΔLISH) domain were cloned by the Gateway® (Invitrogen) method into pDONR221 entry vector and then recombined into a yeast low-copy number CEN destination vector (Addgene, ^70^) to obtain pAG413-promGPD-WDR47 (pSF634), pAG413-promGPD-WDR47-ΔCTLH (pSF636) or pAG413-promGPD-WDR47-ΔLISH (pSF637) plasmids bearing the *HIS3* auxotrophic marker for selection of the transformants on SC-His medium. Plasmid sequences were verified (GATC Biotech). The promCUP1-mCherry-V5-ATG8 (pFL78, *LEU2* selection marker) plasmid was a kind gift from Fulvio Reggiori (Aarhus Institute of Advanced Studies, Aarhus University, Denmark). The *Saccharomyces cerevisiae* wild-type BY4742 (*MATα leu2*Δ*0 ura3*Δ*0 his3*Δ*0 lys2*Δ*0*) reference strain and the *oxa1Δ* (*MATα leu2*Δ*0 ura3*Δ*0 his3*Δ*0 lys2*Δ*0 oxa1::KanMX*) mutant strain were used. The indicated yeast strains were grown at 30°C in rich medium YPD: 1% yeast extract, 2% peptone, 2% glucose, in synthetic complete medium to maintain the plasmid SC-His: 0.67% yeast nitrogen base (YNB, MP Biomedicals) without amino acids, 2% glucose and the appropriate –His dropout mix (CSM-His, MP Biomedicals) or in autophagy induction medium SD-N: 0.17% YNB without ammonium sulphate without amino acids, 2% glucose. Yeast cells were transformed using the modified lithium acetate method ^71^.

For western-blot analysis, total yeast protein extracts were obtained by NaOH lysis of 1.5 unit at OD- 600nm of yeast cells, followed by trichloroacetic acid (TCA) precipitation and the pellet was resuspended in 50 μl of 2X Laemmli buffer plus Tris Base. Samples were incubated 5 min at 37°C prior anti-WDR47 western-blot analysis using standard procedures. TCE (2,2,2-Trichloroethanol) staining was used for total protein detection and loading control ^72^. Images were acquired with the ChemiDoc Touch Imaging System (Bio-Rad).

Drop test growth assays were done on the indicated yeast cells grown from SC-His precultures to exponential phase in YPD for 4 hr at 30°C, prior plating on YPD-2% agar medium as 7 μl drops of serial dilutions at OD-600nm of 0.5, 0.1, 0.01 and 0.005. The plates were incubated at 30°C.

Autophagy was analyzed on yeast cells bearing the empty pAG413 plasmid (control) or expressing Wdr47 (FL, ΔCTLH or ΔLISH) constructs transformed with the mCherry-ATG8 expression vector (pFL78).

Cells were grown in SC-His-Leu +CuSO4 (1 mM) medium to induce the expression of mCherry-V5-ATG8, at OD-600nm 0.5-1 cells were collected before washing with autophagy SD-N medium, and incubation in SD-N medium at 30°C for 4 h prior observation by fluorescent microscopy. Observation was performed with 100X/1.45 oil objective (Zeiss) on a fluorescence Axio Observer D1 microscope (Zeiss) using DsRED filter and DIC optics. Images were captured with a CoolSnap HQ2 photometrix camera (Roper Scientific) and treated by ImageJ (Rasband W.S., ImageJ, U. S. National Institutes of Health, Bethesda, Maryland, USA, http://imagej.nih.gov/ij/).

### Primary neuronal culture

Cortices from E15.5 *Wdr47*^tm1b/tm1b^ or WT mouse embryos were dissected in cold PBS supplemented with BSA (3 mg/mL), MgSO4 (1 mM, Sigma), and D-glucose (30 mM, Sigma). They were then dissociated in Neurobasal media containing papain (20U/mL, Worthington) and DNase I (100 μg/mL, Sigma) for 20 min at 37°C with brief vortexing every 6-7 min. Cells were then incubated for 7 min with Neurobasal media containing Ovomucoïde (15 mg/mL, Worthington), and manually triturated 10 times with a 1000 ml pipette in OptiMeM supplemented with D-Glucose (20mM). Cells were then plated at 2 × 10^5^ cells per 24-well plate or 3.5 × 10^5^ cells per 35 mm diameter glass bottom dish (Cellvis, D35-20-1.5h) coated with poly-D-lysine (1 mg/ml, Sigma) overnight at 4°C and cultured for up to 10 days in Neurobasal medium supplemented with B27 (1×), L-glutamine (2 mM), and penicillin (5 units/ml)-streptomycin (50 mg/ml). Half of the media was changed with fresh supplemented Neurobasal media every 3 days.

### *In vitro* cell death experiments and drug treatments

Cultured neurons were magnetofected with 50ng of Scarlet plasmid at DIV1 using NeuroMag (OZ Bioscience) according to the manufacturer’s protocol and treated at DIV2 with either DMSO, QVd-Oph (Sigma - SML0063) (50uM), Necrostatin (Sigma - N9037) (2 μM), Ferrostatin (Sigma - SML0583) (5 μM) or EpoD (Abcam - ab143616) (10nM). Half of the media was changed every 3 days and neurons were imaged once every day from DIV3 to DIV10 using a Zeiss Axio Observer 7 microscope equipped with a 10x/0.45 plan Apochromat objective controlled by zen 3.3 software, with a CMOS sensor camera (Hamamatsu Orca Flash 4.0 LT Plus) and an incubation chamber (37°C, 5% CO_2_).

### RNA extraction, cDNA synthesis and RT–qPCR

Total RNA was extracted from DIV4 or DIV6 primary neuronal cultures using the NucleoSpin RNA purification kit (Macherey-Nagel) and from human fibroblasts using Trizol reagent (Invitrogen). For RT– qPCR, 1ug of total RNA was reverse transcribed with SuperScript IV (Invitrogen) using random hexamers (2µM) and RT-qPCR was performed with the SYBR Green I Master (LifeScience) and specific primers (**Supplementary Table 4**) according to the manufacturer’s instructions using the Roche 480 LightCycler. Actin was used as the housekeeping gene and the relative amount of target mRNAs was determined using standard curve method.

### RNA sequencing

RNA was isolated from DIV6 primary neuronal cultures obtained from embryos of 3 different pregnant mothers, using the NucleoSpin RNA purification kit (Macherey-Nagel) according to the manufacturer’s instructions. RNA-seq libraries were prepared from 600ng of total RNA using the TruSeq® Stranded mRNA Library Prep kit and the TruSeq® RNA Single Indexes kits A and B from Illumina. The library quality and quantity were checked using an Agilent 2100 Bioanalyzer and a Qubit dsDNA HS Assay Kit. The sample concentration was adjusted to 2.8 nM before sequencing (50 bp single end) on a HiSeq 4000 (Illumina) using HiSeq 3000/4000 SR Cluster Kit and HiSeq 3000/4000 SBS Kit (50 cycles) according to the manufacturer’s instructions.

### Read alignment and quality assessment

Reads were preprocessed using cutadapt ^73^ version 1.10 in order to remove adapter, polyA and low-quality sequences (Phred quality score below 20); reads shorter than 40 bases were discarded for further analysis. Reads mapping to rRNA were also discarded (this mapping was performed using bowtie ^74^ version 2.2.8). Reads were then mapped onto the mm10 assembly of mouse genome using STAR ^75^ version 2.5.3a. Gene expression was quantified using htseq-count ^76^ version 0.6.1p1 with annotations from Ensembl release 102. Samples coming from one of the pregnant mothers were discarded from analysis as a higher plating density which in follow-up experiments was shown to effect timing of cell death, was noted for that culture.

### Differential expression and pathway analysis

Differentially expressed genes (DEGs) between mutant and control samples were identified using R v. 3.3.2 ^77^ and DESeq2 ^78^ version 1.16.1. p-values were adjusted for multiple testing using the Benjamini and Hochberg (BH) method ^79^ and genes with an adjusted p-value smaller than 0.05 were considered to be DEGs. Results of the differential expression analysis can be found in **Supplementary Table 2**. Pathway enrichment analysis of up- and down- regulated genes was performed separately using ClusterProfiler v. 4.8.2 (R v. 4.3.1) ^80, 81^ and all genes found to be expressed in the dataset were used as gene universe. Only pathways with minimum size of 10 and maximum size of 250 genes were included in the analysis. Ontologies with a BH-adjusted p-value < 0.05 were considered significant. Highly similar GO terms were replaced by the most significant one using GoSemSim implemented in ClusterProfiler. In case of ties, pathways with identical gene contribution and p-value were manually collapsed in the most representative one. For disease ontology analysis mouse genes were converted to human orthologs and a gene set enrichment analysis (GSEA) was run using Cluster Profiler by ranking the genes by log2FC and p-values were BH-corrected.

### Cell culture and transfections

All cells provided by the cell culture platform of the IGBMC (Strasbourg) are mycoplasma free (PCR test Venorgem) and have not been authenticated. Human fibroblasts and Human embryonic kidney (HEK) 293T cells were cultured in DMEM (1g/L glucose) (GIBCO) supplemented with 10% Fetal Calf Serum (FCS), penicillin 100 UI/mL, streptomycin 100 µg/mL. Human LCLs were cultured in RPMI 1640 (Gibco) supplemented with 10 % Fetal Bovine Serum (FBS), penicillin 100 UI/mL, streptomycin 100 µg/mL. Mouse neuroblastoma N2A (ATCC) cells were cultured in DMEM (1g/L glucose) (GIBCO) supplemented with 5% FCS and Gentamycin 40µg/mL. All cells were kept in a humidified atmosphere containing 5% CO2 at 37°C.

For expression analysis of truncated *WDR47* constructs HEK cells were transfected with different HA tagged Wdr47 constructs using Lipofectamine 2000 (Invitrogen) according to the manufacturer’s protocol. Expression of transfected constructs was analyzed 48h after transfection by immunoblotting. To study the effect of human variants on WDR47 levels, N2A cells were transfected with different HA tagged WDR47 variants using Lipofectamine 2000 (Invitrogen) according to the manufacturer’s protocol. Expression of transfected constructs was analyzed 24h after transfection by immunoblotting. For MG132 experiments, transfected cells were treated with 10µM MG132 8 hours prior to collection.

### Protein extraction and western blot

Proteins from primary neuronal cultures (DIV4 and DIV6), transfected HEK 293T cells or human fibroblasts were extracted as follows: cells were washed 1X with ice cold PBS, scrapped in 1ml PBS and the cell pellet was lysed with RIPA buffer (0.01 M Hepes, 0.15M NaCl, 0.01M EDTA, 2.5mM EGTA, 0.1% Triton X-100, 0.1% SDS, 1% Na deoxycholate, 2% NaF) ) supplemented with EDTA-free protease inhibitors (cOmplete™, Roche) for 30 min. Cells debris was removed by high speed centrifugation at 4°C for 25 min and protein concentration was determined using Bio-Rad Bradford protein assay reagent.

Samples were denatured at 95°C for 10 min in Laemmli buffer (Bio-Rad) with 2% β-mercaptoethanol and the indicated amount of proteins were resolved by SDS–PAGE and transferred onto PVDF membrane (Immobilon-P) (500ng for actin, alpha tubulin and tyrosinated tubulin, 2µg for acetylated tubulin and 10ug for Wdr47). Membranes were blocked in 5% milk in PBS buffer with 0.1% Tween (PBS-T) and incubated overnight at 4°C with the appropriate primary antibodies (**Supplementary Table 3**) in blocking solution. Membranes were washed 3 times in PBS-T, incubated at room temperature for 1 h with HRP-coupled secondary antibodies (Invitrogen) (**Supplementary Table 3**) at 1:10,000 dilution in PBS-T, followed by 3 times PBS-T washes. The revelation was done by chemiluminescence using SuperSignal West PicoPLUS or SuperSignal™ West Femto Chemiluminescent Substrates (Thermo Fischer) and gels were imaged using the Amersham Imager 600. Relative protein expression was quantified using ImageJ software.

### Live mitochondria imaging and analysis

At DIV4, neurons were transfected with a Venus plasmid (50ng) and mitoDs-Red (250ng) by magnetofection. At DIV6, 10 z-stacks of 0.2 µm step size was imaged from the medial part of the axon (∼ 500-800um from soma) using a confocal microscope Spinning Disk CSU-X1 “Nikon”, equipped with a 100x oil-immersion objective (N.A. 1.4) controlled by the Metamorph 7.10 software, with a CMOS sensor camera (Photometrics Prime 95B) and an incubation chamber (37°C, 5% CO_2_). A home-made ImageJ macro that selects the mitochondria based on red channel intensity was used to quantify the number and morphologic characteristics of the mitochondria.

### Lysosome and mitochondria motility

At DIV4 neuronal cultures were magnetofected with either LAMP1-YFP (500ng) or mito-Dsred (250ng) to label lysosomes and mitochondria respectively. Sparsely labelled cultures were selected to allow selection of single axons and determine their orientation unambiguously. The medial part of the axon (∼ 500-800µm from soma) was chosen and images were acquired every 750 ms for 7 minutes for lysosomes and every 1s for 5 minutes for mitochondria using a Leica spinning disk microscope (CSU-W1) equipped with an Adaptative Focus Control (AFC), a 63x oil-immersion objective (NA 1.4), an Orca Flash 4.0 camera and an incubation chamber (37°C, 5% CO_2_). All acquisition settings were set to keep the signal in a dynamic range and laser powers were kept at minimum to prevent photobleacing. Kymographs were obtained from the time-lapse recordings using a custom written ImageJ plugin Kymo ToolBox v.1.01 (https://github.com/fabricecordelieres/IJ-Plugin_KymoToolBox/releases). Mitochondria and lysosomes that have displacement < 6 µm and <10 µm respectively was considered stationary to avoid bias due to stage drift. All dynamic parameters of intracellular transport are from data that were obtained for each condition in 10-15 axons from at least three independent neuronal cultures.

### Measurement of mitochondrial membrane potential

DIV6 neurons were co-labeled with 20nM potentiometric fluorescent probe TMRE (tetramethylrhodamine, ester ethylique, perchlorate) (Sigma) and 50nM MitoTracker Green FM dye (ThermoFisher) at 37°C 5% CO_2_ for 30 min, maintained with 5nM TMRE for 10 min to achieve equilibration and kept in the same media during subsequent live imaging for a maximum of 20 minutes. Images with a z-stack of 0.3 um were acquired using a Leica spinning disk microscope (CSU-W1), equipped with a 63x oil-immersion objective (NA 1.4), an Orca Flash 4.0 camera and an incubation chamber (37°C, 5% CO_2_). To select mitochondria in the neurites, images were segmented based on MitoTracker Green labeling using the Imaris program and TMRE intensity of each mitochondria was analyzed for 500-1000 mitochondria from 3-5 independent experiments.

### Measurement of mitochondrial redox potential

Neurons were transfected with 250 ng of mitochondria targeted roGFP2 plasmid (Grx1-roGFP2) by magnetofection at DIV1 and live imaging of mitochondria in neurites was performed using a confocal microscope (TCS SP8X; Leica) equipped with 63X oil immersion objective and an incubation chamber (37°C, 5% CO_2_) at DIV6. Confocal settings were pixel size 70.81 (1304x1304 pixels), scan speed 400Hz, laser power 405nm 20%, laser power 488nm 30%, emission bandwidth 498-560, line accumulation of 3 and 2X zoom. For each condition, 405 :488 nm ratio of each mitochondria was calculated in 10-15 axons from at least 3 independent neuronal cultures. Increased 405 :488 ratios toward a more oxidized state suggest increased ROS production.

### Image acquisition and analyses after IUE

All experiments were done in at least three independent replicates and analysis were performed blinded to condition. Cell counting and CC analyses were done in at least three different brain slices of at least three different embryos or pups for each condition. After histological examination, only brains with comparative electroporated regions and efficiencies were conserved for quantification.

#### Counting for neuronal migration

Images for migration analysis were acquired in 1024x1024 mode using confocal microscope (TCS SP8X; Leica) at 20X magnification with a z stack of 1,5 μm and analyzed using ImageJ software. For analyses, upper cortical plate, lower cortical plate, intermediate zone and subventricular zone/ventricular zone were identified according to cell density (nuclei staining with DAPI). The total number of GFP-positive cells in the embryonic brain sections was quantified by counting positive cells within a box of fixed size and the percentage of positive cells in each cortical area was calculated.

#### In vivo CC analyses

Images were acquired using a Zeiss Axio Observer 7 microscope equipped with a 10x/0.45 plan Apochromat objective controlled by zen 3.3 software, with a CMOS sensor camera (Hamamatsu Orca Flash 4.0 LT Plus) and analyzed using ImageJ software. For projecting neuron analyses, the mean fluorescence intensity within a box of fixed width covering the whole axonal thickness at the immediate start of the CC was measured and this value was divided by the mean fluorescence intensity in the cortical plate (**Figure 3C**). For axon extension analysis, CC length was divided into 10 equal bins from the immediate start of the CC till the midline and fluorescence intensity in each bin was measured. Intensity in bin1 was considered 100% and intensity in each bin was normalized to bin1 (**Figure 3D**). For axonal midline crossing analyses, the fluorescence within a box of fixed size placed at the contralateral side of the midline of the brain section was measured and divided by the fluorescence at the ipsilateral side of the midline area (**Figure 3E**). For all analysis background fluorescence intensity was subtracted from the measured intensities.

### Statistics

All statistics were calculated using GraphPad Prism 6 (GraphPad) and are represented as mean +/- sd or mean +/- SEM with the exception of Figure 2 which shows the average between data points for simplicity purposes. All statistical tests used and n numbers have been mentioned in figure legends along with the respective data and statistical details are reported in **Supplementary Table 5**. Graphs were generated using GraphPad and images were assembled with Adobe Photoshop 13.0.1 (Adobe Systems).

### Data Availability

All other relevant data included in the article are available from the authors upon request. The following databases and in silico software were used in the study: Human Gene Mutation Databases (http://www.hgmd.cf.ac.uk/ac/introduction.php?lang=english), the single Nucleotide Polymorphism database (http://ftp.ncbi.nih.gov/snp/), genome aggregation database (gnomAD browser v2.1.1 and v3.1.2 (https://gnomad.broadinstitute.org/)), 1000 genomes (https://www.internationalgenome.org/), University of Santa Cruz Genomic Browser (https://genome.ucsc.edu/index.html) , ClinVar (https://www.ncbi.nlm.nih.gov/clinvar/), Mastermind by Genomenom (https://mastermind.genomenon.com/), Leiden Open Variation Database – LOVD (https://www.lovd.nl/), OMIM – Online Mendelian Inheritance in Man (https://www.omim.org/), Integrative Genomics Viewer - IGV Browser (https://software.broadinstitute.org/software/igv/), Polyphen-2 (http://genetics.bwh.harvard.edu/pph2/), Mutation Taster (http://www.mutationtaster.org/), Sorting Intolerant from Tolerant (SIFT, https://sift.bii.a-star.edu.sg/) and Combined Annotation Dependent Depletion (CADD, https://cadd.gs.washington.edu/). The hWDR47 variants have been deposited in LOVD (Leiden Open Variation Database) v3.0 under the accession numbers #0000953686 (p.(Pro650Leu)), #0000445644 ((p.(Lys592Arg)) and #0000445645 (p.(Asp466His)).

## Acknowledgments

This work was supported by grants from INSERM (ATIP-Avenir program, J.D.G.), the Fyssen Foundation (J.D.G.), the French state funds through the Agence Nationale de la Recherche (JCJC CREDO ANR-14- CE13-0008-01 to J.D.G.; JCJC WDR ANR-18-CE12-0009 to B.Y.; ANR-10-IDEX-0002-02 and ANR-10- LABX-0030-INRT to J.D.G.), SFRI-STRAT’US project [ANR 20-SFRI-0012]; and EUR IMCBio [ANR-17-EURE-0023]), INSERM/CNRS (J.D.G. and S.F.), and University of Strasbourg (J.D.G. and S.F.). E.B. was supported by INSERM, Fondation Jerome Lejeune and ANR. J.R.A. was supported through the IGBMC PhD program and Fondation pour la recherche médicale. L.T. was supported by INSERM through a training grant. S.C.C. is a senior lecturer at the University of Bourgogne; L.T. is a technician; and M.K. a PhD student supported through the Agence Nationale de la Recherche (ANR-11-PDOC-0029-01 to B.Y.). R.L. and N.S. are supported by Ministère de l’Enseignement Supérieur de la Recherche et de l’Innovation and ERC CoG 2021 TransNeuroFate. P.T. is research assistant at the University of Strasbourg. S.F. is a CNRS investigator. B.Y. and J.D.G. are INSERM investigators. H.S. and M.N was funded by the Japan Society for the Promotion of Science, Grant-in-Aids for Scientific Research (JP20H03641 to H.S; and JP21K06819 to M.N.), Japan Agency for Medical Research and Development (AMED; grant numbers: JP23ek0109549, JP23ek0109674, and JP23ek0109637 to H.S.), and Takeda Science Foundation Specific Research Grants (H.S. and M.N.), and HUSM Grant-in-Aid from Hamamatsu University School of Medicine (H.S. and M.N.).

We thank the Imaging Center of IGBMC (https://ici.igbmc.fr/) part of the IBiSA labeled PIQ-QuESt (https://piq. unistra.fr) and of the national infrastructure France Bioimaging (https://france-bioimaging.org), in particular Elvire Guiot and Erwan Grandgirard, for their assistance in the imaging experiments. We are grateful to the staff of the molecular biology service (in particular, Thierry Lerouge and Paola Rossolillo), of the mouse facilities of the Institut Clinique de la souris (ICS) (in particular Sophie Brignon), of the GenomeEast platform (in particular Celine Keime) and of the IGBMC cell culture service for their involvement in the project. We also thank Oktay Cakil for the qPCR with human fibroblasts and Marie-Christine Fischer, Nina Pigeonneau, Perrine F Kretz and Christel Wagner for their helps with histology, image analysis and genotyping for the Nex and CaMKII neuroanatomy studies. We warmly thank Dr. Julien Courchet and Dr. Marine Lanfranchi (Institut NeuroMyoGene, Lyon, France) for sharing plasmids and their helpful comments and advice for mitochondrial assays, Dr. Clement Charenton (IGBMC, Strasbourg, France) for his help on modeling the Wdr47 variants based on Wdr47 structure and Dr. Fulvio Reggiori (Aarhus University, Aarhus, Denmark) for sharing the mCherry-Atg8 plasmid. We also would like to thank Dr.Frederic Saudou, Dr.Chiara Scaramuzzino, Dr. Sandrine Humbert and Mariacristina Capizzi (Grenoble Institute of Neurosciences, Grenoble, France) for helpful comments on experimental design. We are also grateful to Amélie Piton (IGBMC) and to members of B. Y.’s and J.D.G.’s laboratory for discussion and technical assistance and in particular to Elena Brivio for the help with the analysis of RNA-seq data and Aline Dubos for critical reading of the manuscript.

## Author contributions

E.B. conceived and performed the experiments, coordinated and supervised the study, analyzed the data, performed statistical analysis and wrote the manuscript with contributions from all other authors. P.T. conceived and performed *in utero* electroporation, collected the mouse samples, processed the tissues and did immunostainings, did genotyping and provided technical assistance. J.R.A performed lysosome motility and mitochondrial morphology experiments. S.C.C., M.K. and L.T. did neuroanatomical characterization of Nex^Cre^ and CaMKII^Cre^ KO mice. B.R. did the clonings, western blot, transformation and growth test assay for the yeast experiments. S.F performed analysis of yeast data and contribute to the writing of the manuscript. R.L. did the qPCRs and WBs in neuronal cultures. N.S did WBs in N2A cells. S.M., F.K., S.A.T., M.H., F.M., Z.Y., M.N., F.S.A. followed-up the patients and families, provided the clinical and imaging data and contributed to the generation of whole-exome sequencing, bioinformatics tools and analysis of sequencing data. B.Y. and J.D.G. conceived, coordinated and supervised the study and wrote the manuscript with contributions from all other authors.

