## supplementary information for "Bi-allelic variants in *WDR47* lead to neuronal loss causing a rare neurodevelopmental syndrome with corpus callosum dysgenesis in humans"

*Bayam et al.*

Content:

- **Supplementary Figure 1**: Patients with *WDR47* variants, related to Figure 1.
- **Supplementary Figure 2**: Validation of the mouse models used in the study, related to Figure 2.
- **Supplementary Figure 3**: Loss of *Wdr47* induces massive neuronal death at early postnatal stages, related to Figure 3.
- **Supplementary Figure 4**: Effect of different rescue constructs on neuronal migration in control conditions, related to Figure 4.
- **Supplementary Figure 5**: *Wdr47* loss leads to cell death independently of autophagy induction, related to Figure 5.
- **Supplementary Figure 6**: Loss of *Wdr47* in neurons does not impair intracellular transport or pool of dynamic microtubules, related to Figure 6.
- **Supplementary Figure 7**: Effect of hWDR47 constructs with different human mutations on CC and neuronal survival in control conditions, related to figure 7.
- **Supplementary Table 3**: List of primary and secondary antibodies used in this work.
- **Supplementary Table 4**: List of RT-qPCR primers used in this work.
-

**Supplementary Figure 1: Patients with *WDR47* variants, related to Figure 1.**

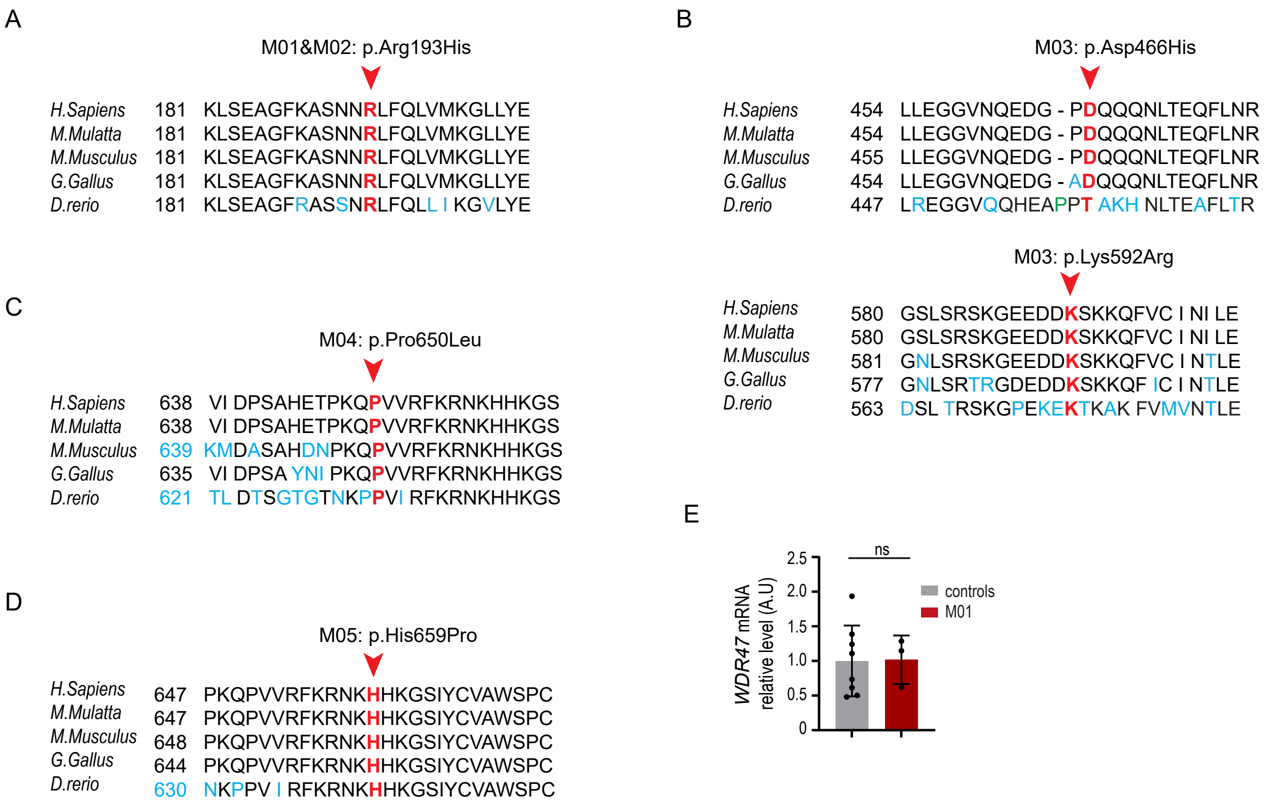


**(A-D)** Alignment of WDR47 protein across several species (human, macaque, mouse, fish, chicken and zebrafish) shows the conservation of mutated amino acid residue (red arrow head) in patients. **(E)** RT-qPCR analyses showing unchanged *WDR47* mRNA levels in M01. Each dot represents one independent measure and data from two individuals used as controls are pooled. Data (means ± s.d) were analyzed by unpaired two-tailed Student t-test, ns, non-significant.

**Supplementary figure 2: Validation of the mouse models used in the study, related to Figure 2.**

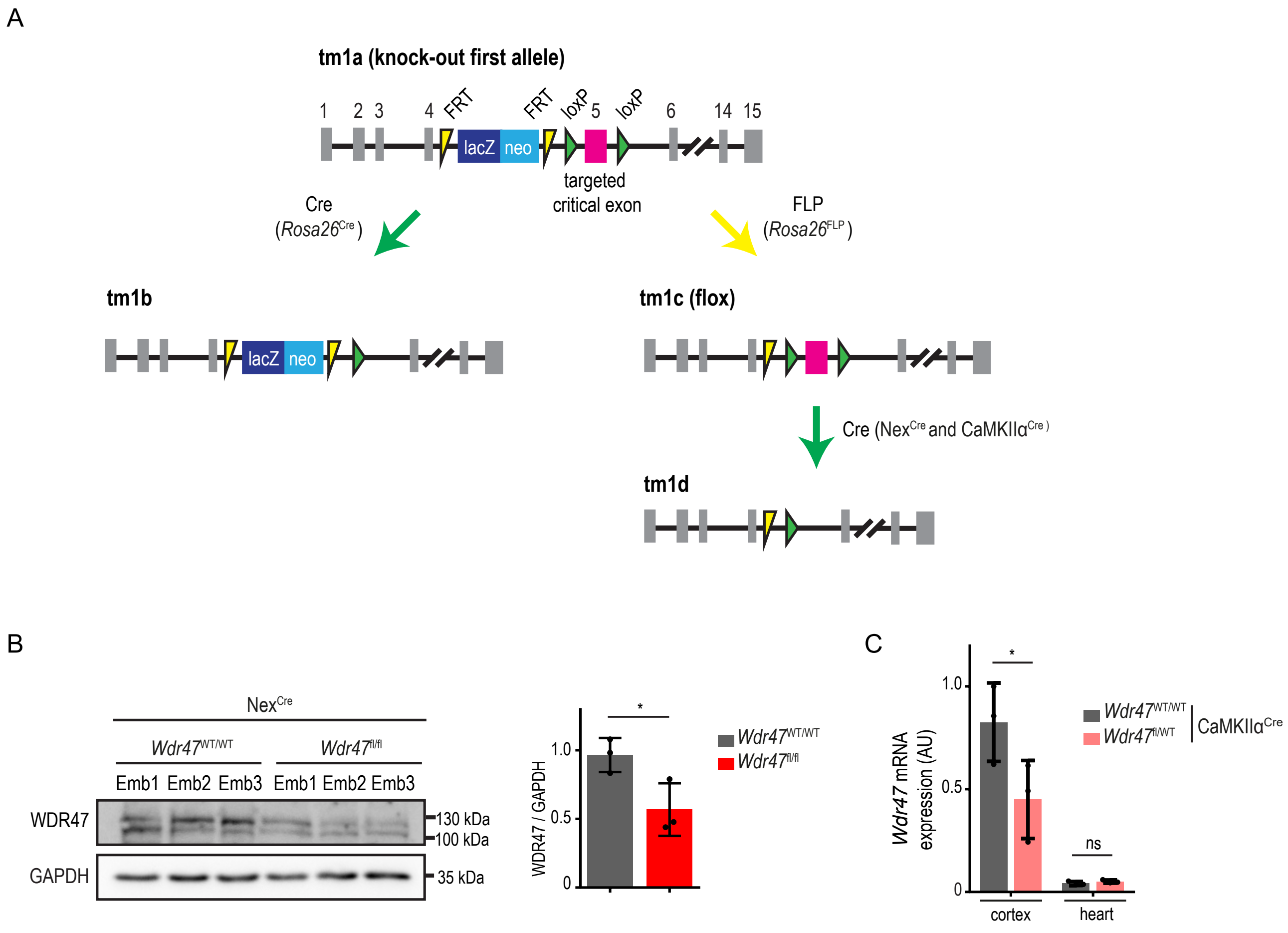


**(A)** Schematic showing allelic construction of the different *Wdr47* mouse models used in the study. FRT, flipase site; LacZ, β-galactosidase gene; loxP, Cre recognition site; neo, neomycin-resistant gene.

**(B)** Western blot analysis of WDR47 in cortical extracts from E18.5 embryos upon deletion of *Wdr47* in early glutamatergic neurons using the $Nex^{Cre}$ mouse line. Gapdh is used as the loading control. Data (means ± sd) from 3 embryos per condition were analyzed by unpaired two-tailed Student t-test. **(C)** RT-qPCR analyses of *Wdr47* mRNA levels in cortical and heart extracts from adult mice upon deletion of *Wdr47* in post-mitotic neurons using the $CaMKII\alpha^{Cre}$ line. Data (means ± s.d) from three samples per condition were analyzed by two-way ANOVA, with Bonferroni's multiple comparisons test. ns, non-significant; $^*P < 0.05$.

**Supplementary Figure 3: Loss of *Wdr47* induces massive neuronal death at early postnatal stages, related to Figure 3.**

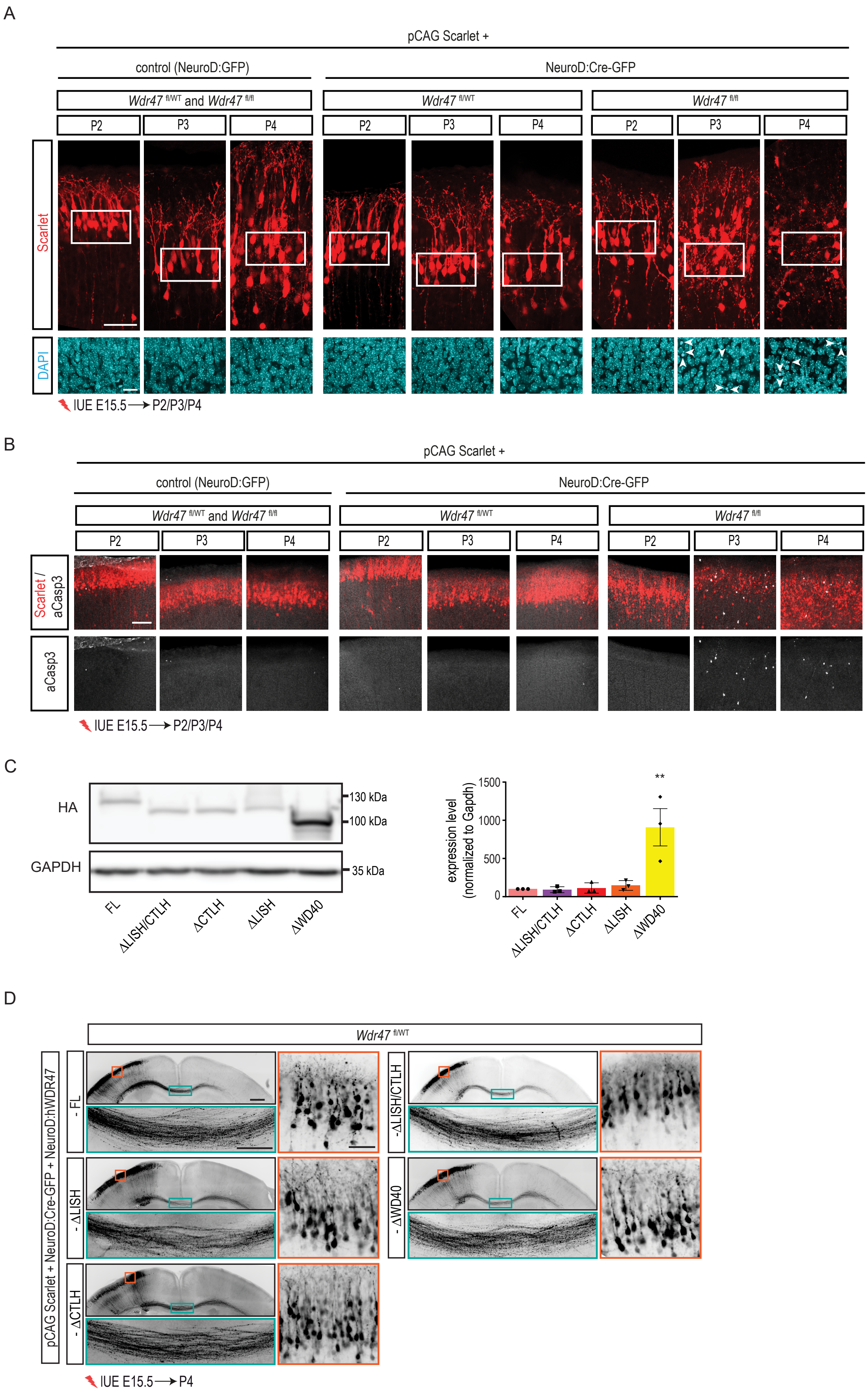

**(A-B)** Coronal sections of P2, P3 and P4 mouse brains electroporated at E15.5 with pCAG:Scarlet plasmid together with either a control (NeuroD:Ires:GFP) or a NeuroD:Cre-GFP vector. Scarlet positive electroporated neurons are depicted in red. **(A)** While *Wdr47*$^{fl/WT}$ neurons and neurons in control conditions keep a proper morphology, *Wdr47*$^{fl/fl}$ neurons lose their bipolar morphology from P3 on. In close-up views of the white boxed area, nucleus is counterstained with DAPI and arrowheads point to pyknotic nuclei. Scale bars: 50µm and 20µm (insets). **(B)** Coronal sections are immunolabelled for activated Caspase3 (aCasp3) (white). Several aCasp3+ cells appear at P3 and P4 in *Wdr47*$^{fl/fl}$ condition. Scale bar: 100µm. **(C)** Western blot analysis of extracts from HEK cells transfected with the indicated HA tagged hWDR47 truncated constructs. Gapdh is used as the loading control. Data (means ± sd) from at least 3 independent experiments. Note that ∆WD40 construct is expressed about 10 times more than the other constructs. **(D)** Effect of different rescue constructs on CC and neuronal survival in control conditions. Coronal sections of P4 *Wdr47*$^{fl/WT}$ mouse brains electroporated at E15.5 with pCAG:Scarlet and NeuroD:Cre-GFP plasmids together with a truncated WDR47 construct. Scarlet positive electroporated neurons are depicted in black. Close-up views of the green and orange boxed area show no effect of the construct on the CC and neuronal survival. Data from at least 3 independent experiments. Scale bars: 500µm, 200µm (green boxed inset) and 50µm (red boxed inset).

**Supplementary Figure 4: Effect of different rescue constructs on neuronal migration in control conditions, related to Figure 4.**

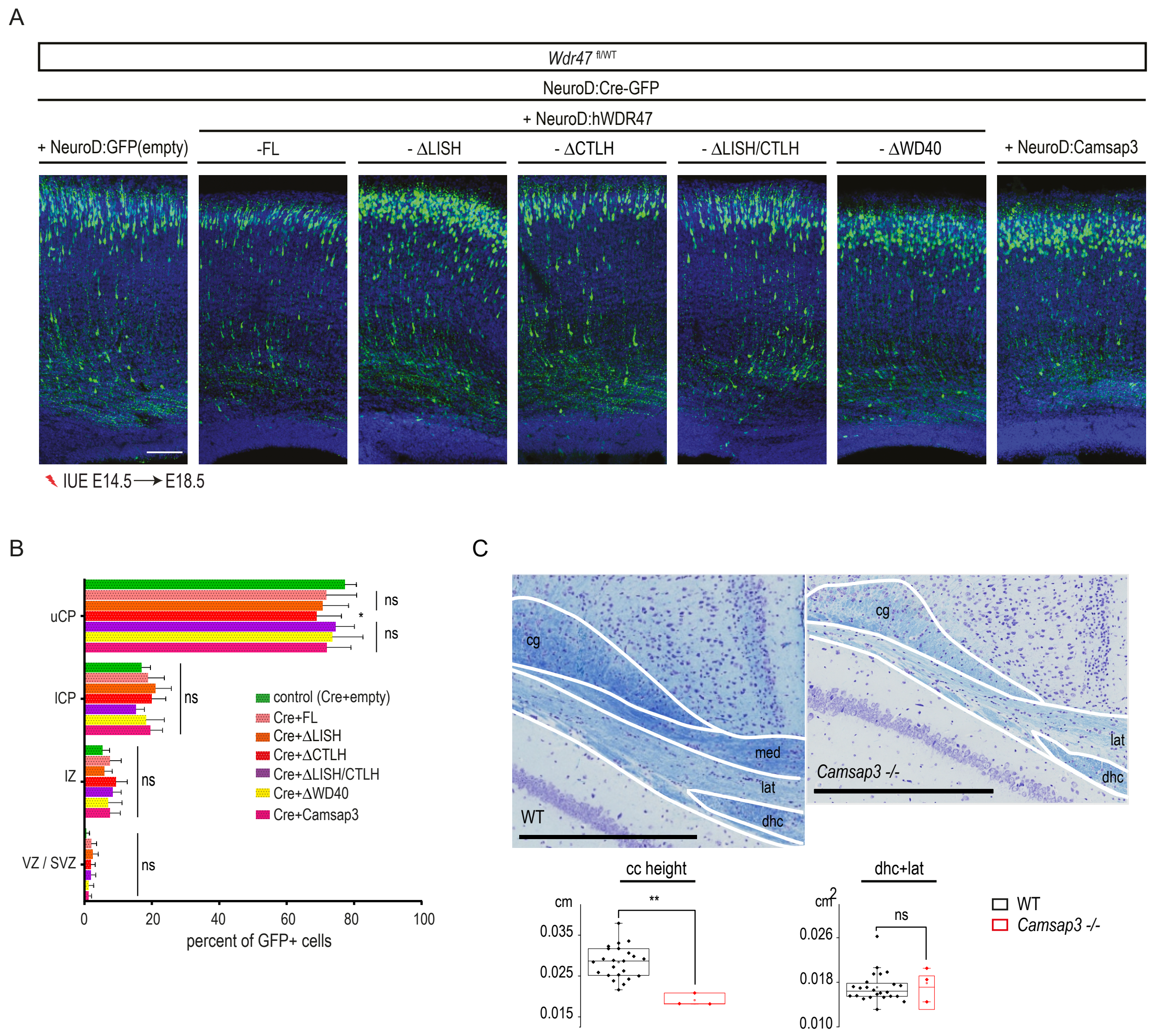


**(A)** Coronal sections of E18.5 *Wdr47*$^{fl/WT}$ mouse cortices four days after *in utero* electroporation with NeuroD-Cre-GFP together with a different NeuroD construct used for rescue experiments. GFP positive electroporated cells are depicted in green. Nuclei are stained with DAPI. Scale bar 100 µm **(B)** Analysis of the percentage of electroporated GFP-cells in different regions (uCP, lCP, IZ and VZ /SVZ) show that apart from the mild effect of ΔWD40 domain construct, none of the constructs have an effect on neuronal migration in control conditions. Data (means ± s.d) from at least three embryos per condition were analyzed by two-way ANOVA, with Bonferroni's multiple comparisons test, ns, non-significant; *P < 0.05. uCP, Upper cortical plate; lCP, Lower cortical plate; IZ, intermediate zone; VZ, ventricular zone; SVZ, subventricular zone. **(C)** *Top.* Representative brain image stained with Nissl-luxol of adult male WT and *Camsap3* KO mice showing the soma of the corpus callosum at Bregma -1.34 mm. *Bottom.* Histogram showing the combined size of the lateral fibers (lat) of the corpus callosum with the hippocampal commissure in 3 male *Camsap3*$^{-/-}$ and 24 matched baseline WT mice of 16 weeks of age bred on a pure genetic background C57BL/6N. Scale bar: 0.05 cm. Data (means ± sd) was analyzed by two-tailed Student's *t*-tests of equal variances. ns, non-significant; $^{**}P < 0.005$. cg, cingulate bundle; dhc, dorsal hippocampal commissure; lat, lateral fibers; med, medial fibers.

**Supplementary Figure 5: *Wdr47* loss leads to cell death independently of autophagy induction, related to Figure 5.**

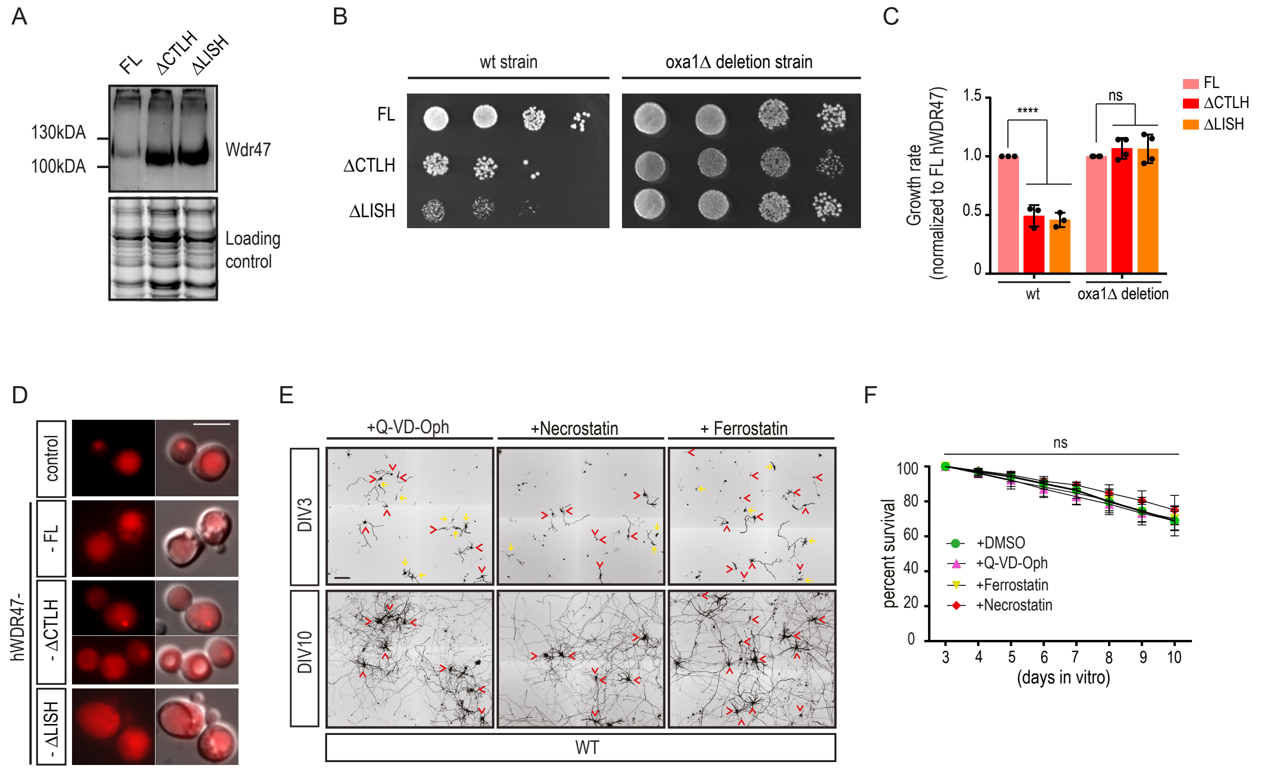


**(A)** WB analysis using yeast extracts showing expression of different hWDR47 constructs (full-length, ΔCTLH and ΔLISH). 2,2,2-Trichloroethanol (TCE) staining is used as loading control. **(B-C)** Yeast drop test growth assay. **(B)** The growth of wt (left panel) and oxa1 mutant (right panel) yeast strains expressing different Wdr47 constructs. **(C)** Quantification of yeast growth expressing different hWDR47 constructs for wt versus oxa1 deletion strains. Note that yeast growth is inhibited upon expression of ΔCTLH and ΔLISH constructs in wt but not in oxa1 deletion strains. Data (means ± sd) from at least 3 independent experiment per condition was analyzed by one-way ANOVA, with Bonferroni's multiple comparison test. **(D)** Fluorescent images of yeast cells, transformed with mCherry-Atg8 plasmid together with or without different WDR47 constructs. Yeast cells were incubated for 4 h in nitrogen starvation medium (SD-N) to induce autophagy. Note that mCherry signal which is strictly localized to the vacuole in control condition becomes diffuse and localizes both to vacuole and cytoplasm upon expression of different WDR47 constructs. Scale bar: 5 µm. **(E-F)** Effect of drugs used for rescue experiments on WT neuronal cultures. **(E)** Representative fields, at DIV3 and DIV10, of WT neuronal cultures treated with Qvd-OPh (50µM) (Caspase inhibitor), Ferrostatin (5 µM) (ferroptosis inhibitor) and Necrostatin (2 µM) (Necroptosis inhibitor) at DIV2. Scarlet positive electroporated neurons are depicted in black. Yellow arrows correspond to neurons that died, red arrowheads correspond to neurons that are alive and could be followed from DIV3 to DIV10. **(F)** Survival of WT neurons from DIV3 to DIV10 upon different drug treatments. Data (means ± sd) from at least 3 independent cultures per condition was analyzed by two-way ANOVA, with Bonferroni's multiple comparison test. ns, non-significant, ****P < 0.0001

**Supplementary Figure 6: Loss of *Wdr47* in neurons does not impair intracellular transport or pool of dynamic microtubules, related to Figure 6.**

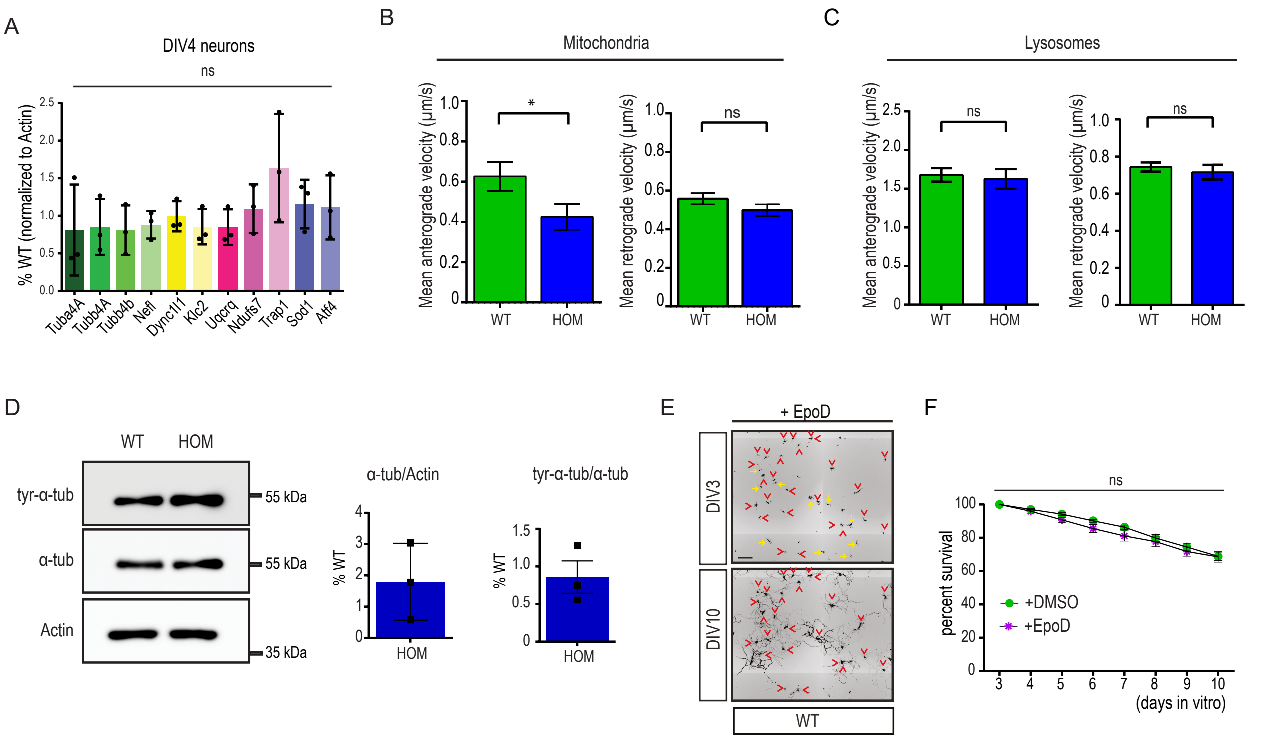


**(A)** RT-qPCR using extracts from DIV4 primary cortical neurons, for the 11 genes that were validated to be differently expressed at DIV6. Data (means ± sd) from at least 3 independent cultures per condition was analyzed by unpaired t-test. **(B-C)** Histograms represent mean anterograde and retrograde velocities of **(B)** mitochondria (anterograde velocity: n=77 for WT and n=36 for HOM; retrograde velocity: n=130 for WT and n=88 for HOM) and **(C)** lysosomes (anterograde velocity: n=216 for WT and n=116 for HOM; retrograde velocity: n=408 for WT and n=173 for HOM). Data (means ± SEM) from at least 3 independent cultures per condition was analyzed by unpaired t-test with Welch correction. **(D)** Western blot analysis showing unchanged levels of alpha tubulin and tyrosinated alpha tubulin in DIV6 HOM (*Wdr47*$^{tm1b/tm1b}$) primary neurons compared to WT. Actin was used as loading control. Data (means ± sd) from 3 independent cultures was analyzed by unpaired t-test. **(E-F)** Effect of EpoD on WT cultures. **(E)** Representative fields, at DIV3 and DIV10, of WT neuronal cultures treated with EpoD at DIV2. Scarlet positive electroporated neurons are depicted in black. Yellow arrows correspond to neurons that died, red arrowheads correspond to neurons that are alive and could be followed from DIV3 to DIV10. **(F)** Survival of WT neurons from DIV3 to DIV10 upon treatment with DMSO and EpoD. Data (means ± sd) from at least 3 cultures per condition was analyzed by two-way ANOVA, with Bonferroni's multiple comparison test. ns, non-significant; *P < 0.05.

**Supplementary Figure 7: Effect of hWDR47 constructs with different human mutations on CC and neuronal survival in control conditions, related to Figure 7.**

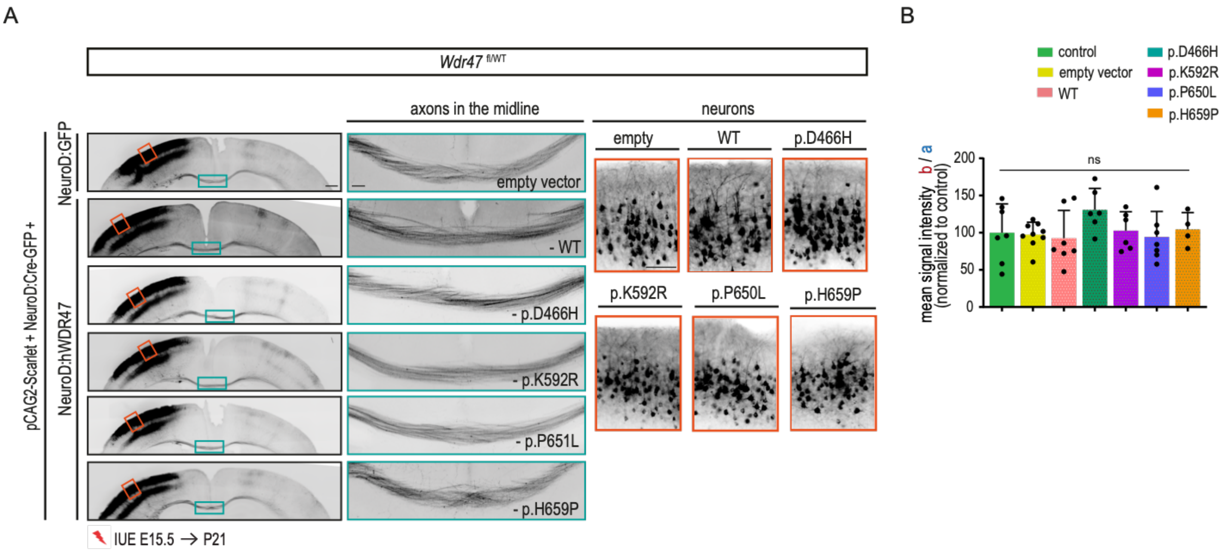


**(A)** Coronal sections of P21 *Wdr47*$^{fl/WT}$ mouse brains electroporated at E15.5 with pCAG:Scarlet and NeuroD:Cre-GFP plasmids and together with either an empty (NeuroD:GFP) or a NeuroD:WDR47 construct with or without the human mutation. Scarlet positive electroporated neurons are depicted in black. Close-up views of the green boxed and red boxed areas show that none of the constructs have an effect on CC or neuronal morphology. Scale bars: 500µm, 100µm (green and red boxed insets). **(B)** CC thickness upon introduction of different hWDR47 constructs. Data (means ± sd) from at least 5 pups per condition were analyzed by one-way ANOVA, with Bonferroni's multiple comparison test. ns, non-significant.

**Supplementary Table 3**: List of primary and secondary antibodies used in this work.

| **$1^0$antibody** | **Host** | **Dilution** | **Used for** | **Company** | **Reference** |
|---|---|---|---|---|---|
| Anti-activated Caspase-3 | Rabbit | 1/100 | IHC | R and D system | AF835 |
| Anti-GFP | Chicken | 1/800 | IHC | Aves lab | GFP-1020 |
| Anti-α-tubulin | Mouse | 1/2000 | WB | Sigma-Aldrich | T9026 |
| Anti-acetylated α-tubulin | Mouse | 1/2000 | WB | Thermo Fisher | 322700 |
| Anti-tyrosinated α-tubulin | Rat | 1/2000 | WB | Sigma-Aldrich | MAB1864-I |
| Anti-β actin coupled HRP | Mouse | 1/40.000 | WB | Sigma-Aldrich | A3854 |
| Anti-HA | Rat | 1:2000 | WB | Roche Life Science products | 11867423001 |
| WDR47 | Rabbit | 1/1000 | WB | Abcam | ab121935 |
| **$2^0$antibody** | **Host** | **Dilution** | **Used for** | **Company** | **Reference** |
| Anti-chicken-488 | donkey | 1/1000 | IHC | Abcam | ab63507 |
| Anti-rabbit-647 | donkey | 1/1000 | IHC | ThermoFisher Sc. | A-31573 |
| Goat-mouse-HRP | Goat | 1/10 000 | WB | ThermoFisher Sc. | G-21040 |
| Goat-rabbit-HRP | Goat | 1/10 000 | WB | ThermoFisher Sc. | G-21234 |
| Goat-rat-HRP | Goat | 1/10 000 | WB | ThermoFisher Sc. | 62-9520 |

IHC, immunohistochemistry, WB, western blot.

**Supplementary Table 4**: List of RT-qPCR primers used in this work.

| Gene | Species | Forward sequence | Reverse sequence |
|---|---|---|---|
| *Actin* | mouse | TATAAAACCCGGCGGC | TCATCCATGGCGAACTGGTG |
| *Dync1I1* | mouse | AGGAAGTGATCGCTCCGGG | CAGACATGTTTGCTTCCTGAATG |
| *Klc2* | mouse | CCGTTCCCAGTGGTGGTATC | CTCCAAGACCAGGCGAAGAG |
| *Ndufs7* | mouse | GTGTCCATGGGGAGCTGTG | CCATAAAGGAGTGCTTCGGC |
| *Nefl* | mouse | CCGGGGTATGAACGAAGCTC | CAGTTTGTTGATTGTGTCCTGC |
| *Sod1* | mouse | GGGAAGCATGGCGATGAAAG | GGTTCACCGCTTGCCTTCTG |
| *Trap1* | mouse | GTCCCGGGTACAAGATGTGG | CATCCAGATGGCCTGCAAAG |
| *Tuba4A* | mouse | CTACGTGAGACGTACAGCCC | TGGTCTTATCGCTGGGCATC |
| *Tubb4A* | mouse | AGAGGAGGCTGAAGAGGAGG | TGGGGTCCTAGGGAATGAGG |
| *Tubb4B* | mouse | TCTACAGCTGTTCCGCAGTC | CTCGTCGCTGATTACCTCCC |
| *Uqcrq* | mouse | TAGGCCGTGGGAGGTTTTTC | AAAGGGCGACAAGCTGTAGG |
| *Atf4* | mouse | GGTGGCCAAGCACTTGAAAC | CGGAAAAGGCATCCTCCTTG |
| *WDR47* | human | TCCTGAGGCAGCTAATACTTG | TCGCGTTGTTAACACATAAAGC |
